# High-precision mapping of nuclear pore-chromatin interactions reveals new principles of genome organization at the nuclear envelope

**DOI:** 10.1101/2021.05.10.443506

**Authors:** Swati Tyagi, Juliana S. Capitanio, Jiawei Xu, Fei Chen, Rahul Sharma, Jialiang Huang, Martin W. Hetzer

**Affiliations:** The Salk Institute for Biological Studies, La Jolla, CA, USA; Paul F. Glenn Center for Biology of Aging Research at the Salk Institute, La Jolla, CA, USA; School of Biological Sciences, University of California San Diego, La Jolla, CA 92093, USA; State Key Laboratory of Cellular Stress Biology, School of Life Sciences, Faculty of Medicine and Life Sciences, Xiamen University, Xiamen, Fujian 361102, China; Institute of Science and Technology Austria, 3400 Klosterneuburg, Austria

## Abstract

The role of nuclear pore complexes (NPCs) in genome organization remains poorly characterized due to technical limitations in probing genome-wide protein-DNA interactions specific to the nuclear periphery. Here, we developed a new sensitive method, NPC-DamID, which combines *in vitro* reconstitution of nuclear import and DamID technology. The fixation-free method identifies chromatin interactions at the NPCs in intact nuclei from cells and tissues. We found that NPCs are preferentially associated with common and hierarchically arranged super-enhancers (SEs) across multiple cell types. We also uncovered phase-separated condensates at NPCs that compartmentalize and concentrate transcriptional coactivators and structural proteins at SE-regulated genes. Our results support NPCs as anchoring sites for SE regulatory hubs and cell-type-specific transcriptional control.

## INTRODUCTION

The spatial organization of the genome in a mammalian nucleus is critical for the transcriptional output of a cell which in turn is essential for cell fate determination and function ^1^. Nuclear landmarks such as the nuclear envelope (NE) play important roles in orchestrating genome architecture and transcription regulation. For example, the interaction between chromatin and the lamina at the NE forms hundreds of large Lamin-associated domains (LADs) of genomic DNA, which are primarily heterochromatin-containing genes with typically low expression ^2^. While the association between lamina and DNA is well understood, the specific chromatin interaction sites of nuclear pore complexes (NPCs), which collectively occupy approximately 20% of the area of NEs, have not been adequately mapped.

NPCs are megadalton protein complexes, composed of ∼30 different subunits called nucleoporins (Nups), that regulate the transport of cargo molecules between the nucleus and cytoplasm ^3^. Recent findings have uncovered additional roles for NPCs and individual Nups in gene regulation and genome organization ^3–5^. However, our functional understanding of NPC-genome interactions has been limited in terms of spatial and temporal resolution and restricted to few cell types due to technical challenges in probing the dynamic association of membrane-bound proteins with chromatin through antibody-based methods like ChIP-seq ^6–8^. For example, the efficiency of ChIP-seq is lower for membrane-embedded proteins like Nups due to the poor accessibility of epitopes of NPC proteins to the antibodies. Moreover, the dynamic interactions between Nups and chromatin are difficult to capture through ChIP-seq. As an alternative, *in vivo* DamID for Nups has been the method of choice to avoid using antibodies for studying NPC-chromatin interactions. The *in vivo* DamID-seq method is independent of antibody usage because transgenic cell lines are used in which Dam-fused proteins are expressed at typically very low levels ^9^. Dam is an enzyme from *E. coli* that, when fused to a protein of interest, positions an orthogonal methylation modification to adenines at GATC of interacting genomic DNA ^9^. The methylated DNA is then sequenced to identify the genomic DNA associated with the protein of interest ^10^. In this way, DamID-seq has been successfully applied to many membrane-embedded proteins, including Nups ^11–13^. However, the expression of Dam fused Nups for DamID-seq experiments in cell lines poses difficulties such as over-expression artifacts, possible toxicity of Dam fused proteins, and loss of temporal resolution due to cumulative DNA methylation and the need for a long time of expression (24-36 h) of Dam fused proteins ^7^. Moreover, Nups like Nup153 or Nup98, which have been previously targeted to study NPC-chromatin associations, are mobile Nups (i.e., they also can bind chromatin at sites that are not associated with the NPCs) ^11, 13–15^. Hence, it is challenging to distinguish genomic DNA interactions at the nuclear periphery from those in the nucleoplasm in these studies.

To overcome some of these limitations of conventional DamID-seq experiments, we have developed a complementary method called NPC-DamID, explicitly designed to identify NPC-genome interactions at the nuclear periphery. Two different assays have been adapted to develop NPC-DamID. The well-established *in vitro* reconstitution of nuclear transport assay (semi-permeabilized cell assay) is combined with DamID-seq to target the Dam fused nuclear transport receptor Importin beta (Impβ) to NPCs (Fig. 1a). This method bridges cell biological and genomic studies and remains independent of the use of an antibody or the over-expression artifacts of Dam fused protein. Since there is no requirement for the generation of transgenic cells, this method can be applied to multiple unfixed cells and tissue types from different species. Here, we demonstrate the use of NPC-DamID to understand the features and mechanisms behind NPCs as genome organizers.

**Figure 1.**
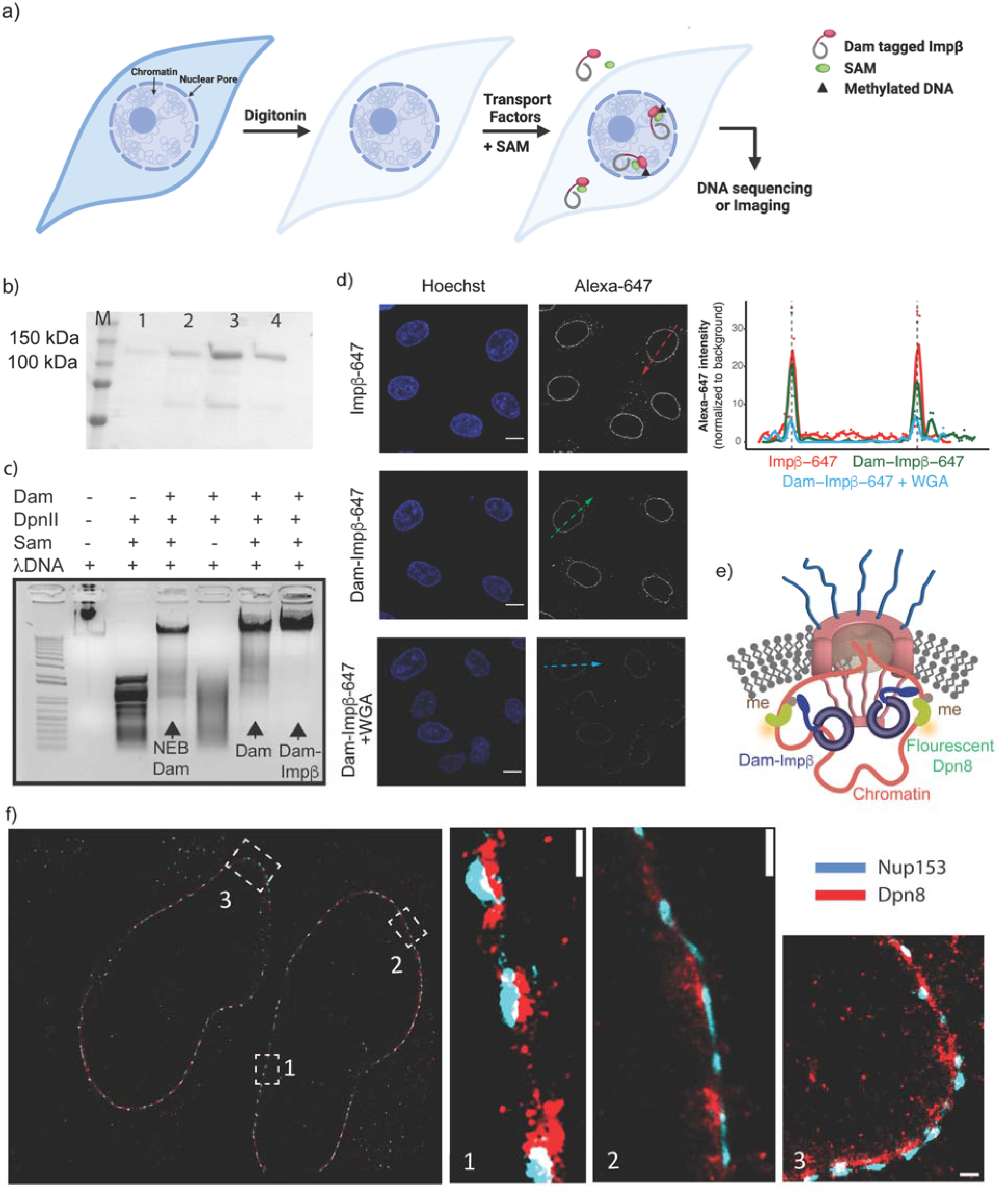
NPC-DamID, a new method to probe NPC-genome interactions at the nuclear periphery. a) Schematic representation of *in vitro* DamID. The cells are semi-permeabilized with a low concentration of digitonin (0.001-0.0025% w/v) so the plasma membrane is preferentially permeabilized, leaving the nuclear membrane intact. Dam-fused nuclear import factor Dam-Impβ is supplied to the cells along with the other transport factors and SAM (s-adenosyl methionine), efficiently targeting Dam-Impβ to the NPCs and subsequently methylating NPC-associated chromatin. In the next steps, cells were lysed to extract genomic DNA to prepare the DamID-seq libraries. b) Coomassie gel of recombinantly expressed and purified Dam-Impβ (molecular weight 132 kDa) through size exclusion chromatography (lane 1-4), lane M is protein marker. c) EtBr-stained agarose gel showing the enzymatic activity of Dam-Impβ. d) Targeting of fluorescently labeled Dam-Impβ and wt Impβ labeled with Alexa 647 to NPCs in HeLa cells after semi-permeabilization and NPC-DamID assay. WGA was added to the assay to block the NPCs. The lack of nuclear rim signal for labeled Dam-Impβ in the presence of WGA shows specific localization of Dam-Impβ at the periphery through NPCs. The scale bar is 10 µm. The arrows indicate the planes used for line plots. The dashed lines in the line plot indicate the limits of the nuclear peripheral. e) Schematic representation of Dam-Impβ methylated DNA imaging in the vicinity of NPCs using fluorescently labeled Dpn8. f) dSTORM images of IMR90 cells treated with Dpn8 after NPC-DamID assay. The cells show co-staining of Dpn8 (staining for NPC-interacting genomic regions) and Nup153 (marking NPCs). The scale bar is 500 nm.

We have recently identified an unexpected link between Nups and super-enhancers (SEs) ^11^. SEs are the cluster of enhancers in close genomic proximity with unusually high levels of transcription factors and activators ^16^. SEs regulate the activity of genes important for cell-type specification ^16^. Research from other groups has also shown similar associations of Nups with SEs by targeting a specific Nup either through DamID-seq or ChIP-seq experiments ^17, 18^. However, since individual Nups are also present inside the nucleus and can interact with chromatin in the nucleoplasm ^3^, no detailed maps exist that precisely and reliably identify NPC-genome interactions at the NE. This results in a lack of understanding of the molecular mechanisms behind these associations and their consequences on cell fate determination and genome organization.

In this study, we mapped NPC-SE interactions across multiple cell types. We uncovered unique features for SEs specifically located at the nuclear periphery, furthering our understanding of the importance and role of NPCs in the function and organization of NPC-SEs.

## RESULTS

### NPC-DamID combines *in vitro* nuclear import assay with DamID to probe NPC-genome interactions at the nuclear periphery

To overcome the challenges associated with *in vivo* DamID-seq and probe the specific chromatin association of NPCs at the nuclear periphery, we developed a method called NPC-DamID. NPC-DamID uses targeting of recombinantly expressed and purified Dam-fused nuclear transport receptor Importin beta (Impβ) to NPCs in a reconstituted nuclear import assay (Fig. 1a, b, Supplementary Fig. 1 a-d). Since Impβ has a high affinity to NPCs, we predicted that Dam fused Impβ (Dam-Impβ) would stably bind to NPCs embedded in the NE when the plasma membrane of cells was permeabilized with low concentration digitonin (Fig. 1a), resulting in specific methylation of DNA in the vicinity of NPCs.

Before reconstituting the nuclear import assay for NPC-DamID, we tested the *in vitro* methylation activity of recombinantly expressed and purified Dam and Dam-Impβ. We incubated unmethylated lambda DNA with these proteins at room temperature (RT) for 30 min. We then digested the DNA with DpnII, a restriction enzyme that can only cleave unmethylated GATC DNA sequences. We found that purified Dam and Dam-Impβ can efficiently methylate lambda DNA in 30 min, similar to commercially available Dam (NEB Dam) (Fig. 1c, Supplementary Fig. 1e).

Next, we confirmed the proper localization of Dam-Impβ at the nuclear periphery by imaging the fluorescently labeled protein after performing the NPC-DamID assay. We observed a nuclear rim signal for Dam-Impβ labeled with Alexa 647 similar to recombinantly expressed purified and fluorescently labeled wt Impβ (Fig. 1d, Supplementary Fig. 1d). However, in the presence of wheat germ agglutin (WGA), a small molecule that blocks NPCs, Dam-Impβ was unable to bind to NPCs, thus serving as a negative control (Fig. 1d). We also confirmed the active import activity of Dam-Impβ, which is dependent on RanGTP through the localization of nuclear localization signal (NLS) containing cargo inside the nucleus (Supplementary Fig. 1f).

Next, we wanted to test if Dam-Impβ can specifically label chromatin in the vicinity of NPCs and if this localized DNA methylation can be visualized microscopically. To this end, we exploited a known fragment of the Dpn1 enzyme, which binds to GA^me^TC residues without cleaving the DNA ^22^(Fig. 1e). Previously, a similar fragment had been used as a tracer for *in vivo* DamID experiments for imaging lamina-associated domains (LADs), which are large megabase pair long stretches of DNA sequences and thus relatively easy to visualize ^23^. We expressed and purified the recombinant fragment (Dpn8) fused with either flag or GFP tag (Supplementary Fig. 1g-j). We added Flag-Dpn8 to the semi-permeabilized cells after washing off the reagents from the NPC-DamID assay. Subsequently, through confocal imaging, we observed that Flag-Dpn8 labels the entire nuclear volume when cells were incubated with Dam alone, indicating that Dam diffuses freely through NPCs during the assay and places methylation marks across the entire genome (Supplementary Fig. 1k). When Flag-Dpn8 was used to label cells treated with Dam-Impβ, only a nuclear peripheral signal was observed, similar to a cell line expressing lamin B-receptor fused to Dam (Dam-LBR). This shows the selective methylation of genomic DNA associated with NPCs through Dam-Impβ. Further, this peripheral signal was lost in the presence of WGA, which prevents Dam-Impβ from binding NPCs, and in the presence of Impβ alone, which cannot methylate DNA (Supplementary Fig. 1k). To determine the working distance range of the NPC-DamID assay, we performed dual-color super-resolution microscopy (dSTORM). We were able to visualize individual NPCs labeled with Nup153 antibody-Alexa 488 and associated genomic DNA labeled with Dpn8-Alexa647, which confirms the selective targeting of nuclear peripheral genomic DNA through the NPC-DamID method (Fig. 1f). We observed that most NPCs uniformly had a signal for associated genomic DNA, and methylation pattern ranges from 200-600 nm from the nuclear periphery. These results show that NPC-DamID provides a method for high-resolution imaging of NPC-associated genomic DNA at the nuclear periphery.

### NPC-DamID sequencing allows the identification of chromatin associated with NPCs and eliminates the nucleoplasmic signal from mobile Nups

To understand the NPC-mediated genome organization at the NE, we must determine the specific genomic regions associated with the NPCs. As in the *in vivo* DamID method, after performing the NPC-DamID assay, we enrich GA^me^TC methylated DNA fragments from the genomic DNA to prepare libraries for DNA sequencing (see Methods section). We used Dam alone for background correction of the sequencing data for Dam-Impβ. We determined the active enzymatic units for purified Dam and Dam-Impβ through an *in vitro* lambda DNA protection assay against the DpnII enzyme (see Methods). One enzyme unit is defined as the concentration of Dam or Dam-Impβ required to completely methylate 1 µg of lambda DNA at room temperature (RT) in 30 minutes, protecting the DNA from DpnII mediated cleavage (Supplementary Fig. 2a-b). In the NPC-DamID assay, the same active enzyme units for Dam and Dam-Impβ were used to ensure unbiased normalization of the data between experiments. The kinetic assay for Dam and Dam-Impβ revealed that 5 units of enzyme protected 2 µg of lambda DNA in 20 min (Supplementary Fig. 2c-d). Thus, we used 5 units of Dam or Dam-Impβ in the nuclear transport assay to target proteins in semi-permeabilized cells for 20 minutes at RT. Our results from sequencing experiments show clear peaks for Dam-Impβ, which are significantly enriched over Dam alone and wt Impβ (Fig. 2a). Dam-Impβ peaks are also significantly non-overlapping with the LADs, indicating the assay’s specificity (Fig. 2a, Supplementary Fig. 2e). To further assess the efficiency of the assay, we added the NPC-blocking small molecule WGA and found no detectable peaks above the background control of Dam alone (Fig. 2a, Supplementary Fig. 2f). This shows that the semi-permeabilization left the NE intact; hence, the peaks for Dam-Impβ are selective for Dam-Impβ localized at NPCs.

**Figure 2.**
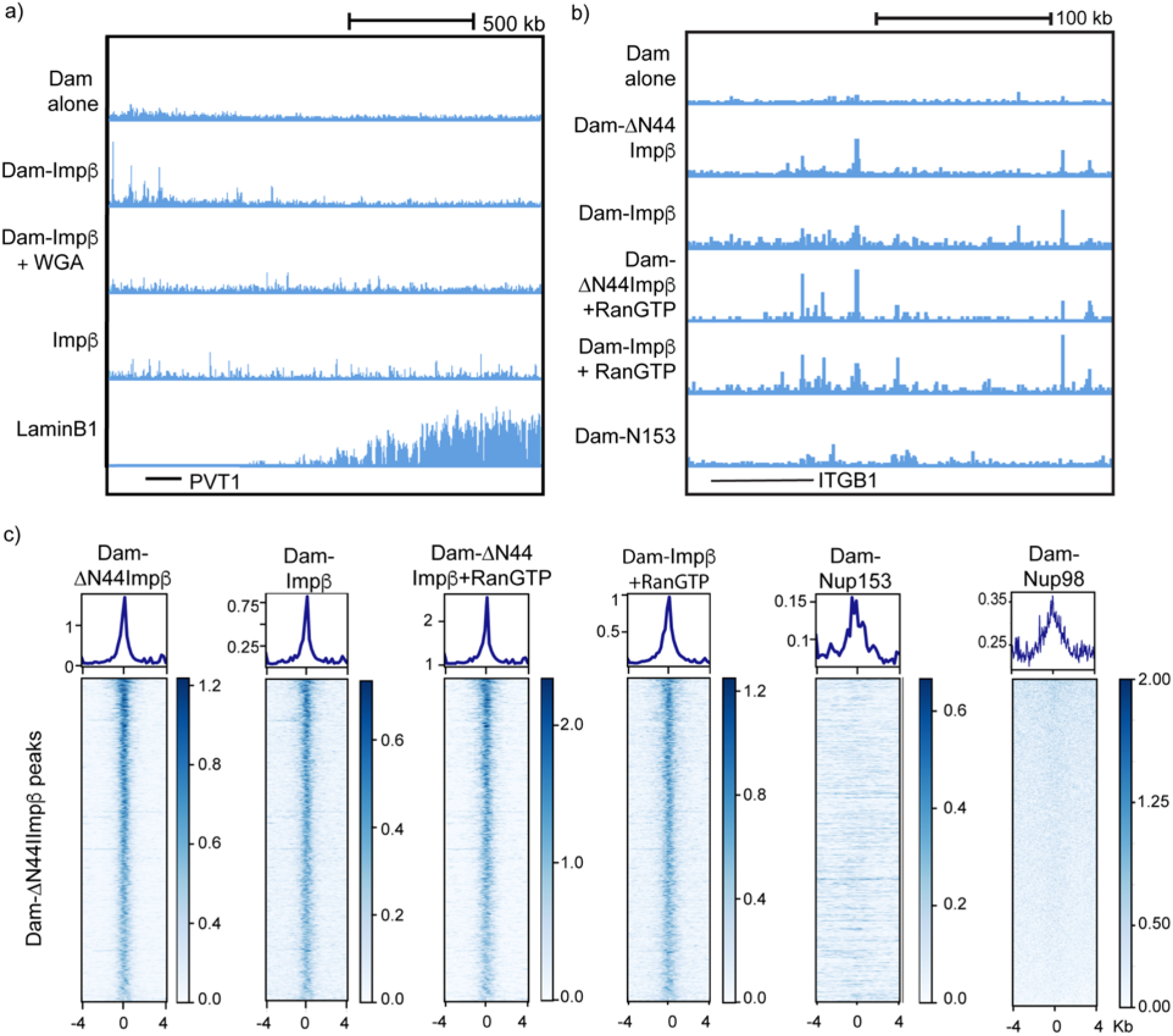
NPC-DamID is a robust and reproducible method to study NPC interaction with chromatin. a) Normalized NPC-DamID DNA sequencing tracks for Dam alone, Dam-Impβ, Dam-Impβ with WGA, wt Impβ, and Lamin B1 for the genomic region around the PVT1 gene in HeLa cells. b) Normalized NPC-DamID DNA sequencing tracks for Dam alone, Dam-ΔN44Impβ, Dam-Impβ, Dam-ΔN44Impβ with RanGTP, Dam-Impβ with RanGTP, and Dam-Nup153 for the genomic region around the ITGB1 gene in IMR90 cells. c) Sequencing reads distribution centered around Dam-ΔN44Impβ peaks for Dam-ΔN44Impβ, Dam-Impβ, Dam-ΔN44Impβ+ RanGTP, Dam-Impβ+RanGTP, Dam-Nup153 for IMR90 and NPC-bound Dam-Nup98 in HeLa cells.

Impβ is known to transiently move from NPCs at the nucleoplasmic side while releasing cargo, a step dependent on the RanGTP binding property of Impβ ^24^. This raises the possibility that Dam-Impβ might be released from the NPCs during the reaction, methylating chromatin regions in the nucleoplasm. To reduce the impact of such diffusive molecules of Dam-Impβ on the sequencing results, we expressed a dominant mutant of Impβ fused to Dam, Dam-ΔN44 Impβ. This mutant does not bind to RanGTP and therefore blocks nuclear import activity by stably binding to the nuclear face of NPCs ^25^. When sequencing data were compared for Dam-Impβ and Dam-ΔN44 Impβ, we found similar peaks (78% overlap) in both data sets, with the mutant Dam-ΔN44 Impβ showing lower background noise in the data (Fig. 2b, Supplementary Fig. 2f). We observed that the presence of RanGTP also did not significantly influence either Dam-ΔN44 Impβ or Dam-Impβ peaks (Fig. 2b) ^24^. For all the assays in this study, we have used Dam-ΔN44 Impβ for all the experiments due to the lower background, unless otherwise mentioned.

Next, we compared NPC-DamID with *in vivo* DamID-seq data previously obtained using Dam-Nup153 expressing IMR90 cell line (Fig. 2c, Supplementary Fig. 2g) or Dam-Nup98 expressing HeLa cells (Fig. 2c). While there is a significant overlap between the two Dam-Nup data sets and their respective NPC-DamID peaks, many additional peaks in Dam-Nup data do not overlap with NPC-DamID data. These additional peaks are likely the result of the mobile nucleoplasmic fraction of Nup153 or Nup98 binding to chromatin. To test this idea, we performed three-dimensional genome modeling using Chrom3D ^26^ to simulate either the position of NPC-DamID peaks or Dam-Nup153 peaks relative to known LADs sequences at the nuclear periphery (Supplementary Fig. 2h-i). The results show that the distance distribution of Dam-Nup153 peaks from the nuclear periphery is bimodal. The fraction of peaks closer to the nuclear periphery overlaps with NPC-DamID peaks, while the other fraction is farther away from the NE, likely representing the nucleoplasmic interaction between Nup153 and chromatin. In line with this idea, we compared the overlap between NPC-DamID and cut&run for Nup93, a nucleoporin that does not bind chromatin in the nucleoplasm. Despite the previously mentioned methodological issues of utilizing an antibody-based method to detect the interaction of Nup93 with chromatin, we still observed an increase in the overlap of NPC-DamID peaks with Nup93 versus Nup153 in IMR90 cells (Supplementary Fig. 2g).

Lastly, we have used fluorescent in situ hybridization (DNA-FISH) to validate the peripheral localization of NPC-interacting loci. As observed for LADs, NPC-interacting loci are more frequently localized in the peripheral zone of the nucleus, adjacent to the nuclear pores embedded in the nuclear envelope (Supplementary Fig. 2j). In all, NPC-DamID is a robust method that specifically methylates NPC-associated genomic regions.

### NPC-associated genomic regions are enriched in enhancer regulatory elements and actively expressed genes

After having successfully established and characterized the NPC-DamID method, we were interested in understanding the chromatin regions associated with NPCs. To this end, we incubated Dam-Impβ with different semi-permeabilized (i.e. unfixed) cells, including HeLa, IMR90, and mouse C2C12 myoblasts. When we analyzed the distribution of varying histone modification ChIP-seq reads centered at NPC-DamID peaks, we found that the NPC-DamID peaks preferentially enrich at open chromatin marks like H3K27Ac and H3K4Me3 as opposed to a repressive mark like H3K9Me3 (Fig. 3a). We further tested functional features of these genomic regions by comparing them to the 18 chromatin states of the ChromHMM model for IMR90 ^27^. Consistent with the enrichment of enhancer and promoter histone marks, we observed the most significant overlap with active enhancers among the 18 different chromatin states for NPC-DamID peaks (Fig. 3b). The next significant overlap was found with weak enhancers and weak transcription regions, followed by transcription start sites (TSS). To check the transcriptional output of NPC-DamID peaks in IMR90 cells, we compared the transcription of NPC-associated genes with a randomly chosen equal number of other genes. We have found that the average transcriptional output from NPC-interacting genes is higher than that of random genes (Fig. 3c, Table S1). This suggests that NPC-associated genomic regions are primarily actively transcribed chromatin regions. These features of NPC-interacting chromatin were found to be consistent in the human HeLa and mouse C2C12 cell lines (data not shown).

**Figure 3.**
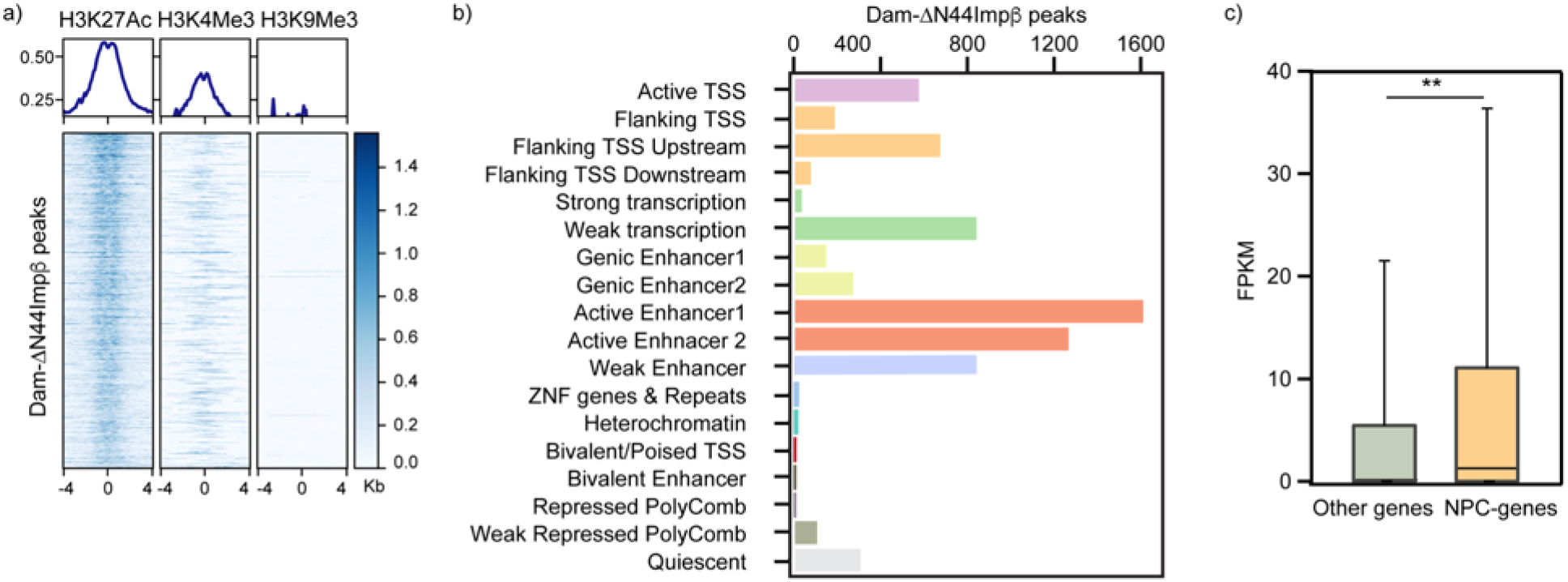
Genomic DNA interacting with NPCs contains mainly regulatory elements. a) Distribution of H3K27Ac, H3K4Me3, H3K9Me3 ChIP-seq reads centered around NPC-DamID peaks (Dam-ΔN44Impβ) in IMR90 cells. b) Percentage overlap of NPC-DamID peaks with the genome-wide annotation of IMR90 from the 18-state ChromHMM model. c) Transcriptional output comparison for genes associated with NPCs versus the same number of randomly selected other genes (NPC-bound genes are shown in Table S1, ** p< 0.01).

Since NPC-DamID does not require genetically modified cells, it also applies to tissue samples. Thus, we could map chromatin associations with NPCs in unfixed human pancreatic islet tissues (which would have been challenging with *in vivo* DamID or ChIP-seq). After dissociating islets into a single-cell suspension, we performed the assay using about 50,000 cells. Interestingly, even though this represents a pool of multiple cell types found within islets, we still observed that NPC-interacting chromatin was enriched in histone marks for enhancer (H3K27Ac) and promoter (H3K4Me3) regions (Supplementary Fig. 3), similar to other cell types tested. The easy applicability of this method to complex tissue samples highlights a significant advantage of NPC-DamID over other methods of NPC-chromatin interaction detection. This way, NPC-DamID allows us to expand our understanding of the genomic associations at the periphery from cells to tissues.

Taken together, across multiple cell types, NPCs interact with open chromatin regions enriched in enhancers and promoters, and the NPC-associated genes show high transcriptional output.

### NPCs associate with super-enhancer regions in different cell types

While the association between Nups like Nup153, Nup93, Nup133, and Nup98 and super-enhancers (SEs) has been established^11, 17, 18^, we wanted to generate high-precision maps of these interactions in multiple cell types to identify SEs that are specifically associated with NPCs at the NE. To do this we performed NPC-DamID sequencing for HeLa, IMR90, and C2C12 myotubes. We found a consistent and significant overlap (40-65%) between SEs and NPC-DamID peaks for all cell types (Fig. 4a). To further understand the role and importance of these interactions, we investigated how NPC-associated SEs (NPC-SEs) differ from SEs that do not associate with NPCs (Table S2). Although SEs are highly specific to a cell type regulating genes crucial for cell-fate determination, recently, it has been shown that some SEs can be shared between two or more cell types ^19^. We wanted to investigate if NPC-SEs are specifically enriched in these common SEs compared to non-NPC-SEs. We generated a cell type consensus score for NPC-SEs, non-NPC-SEs, and all SEs identified in IMR90 and Hela cells across ten different cell types. Our results show that NPC-SEs have a slightly higher consensus score than non-NPC-SEs, indicating that NPC-SEs have higher enrichment for common super-enhancers (Fig. 4b, Supplementary Fig. 4a). These results suggest that SEs associated with NPCs are relatively more shared between different cell types, and the genes regulated by these SEs are responsible for essential cellular functions needed for cell differentiation (Supplementary Fig. 4b). Common SEs are strongly associated with fast recovering chromatin loops after sequential cohesin depletion and restoration ^19^. This leads us to speculate on the intriguing possibility that NPC-linked SEs could play a role in establishing chromatin organization by facilitating the formation of early higher-order chromatin loops that later become scaffolds for cell-type-specific loop formation.

**Figure 4.**
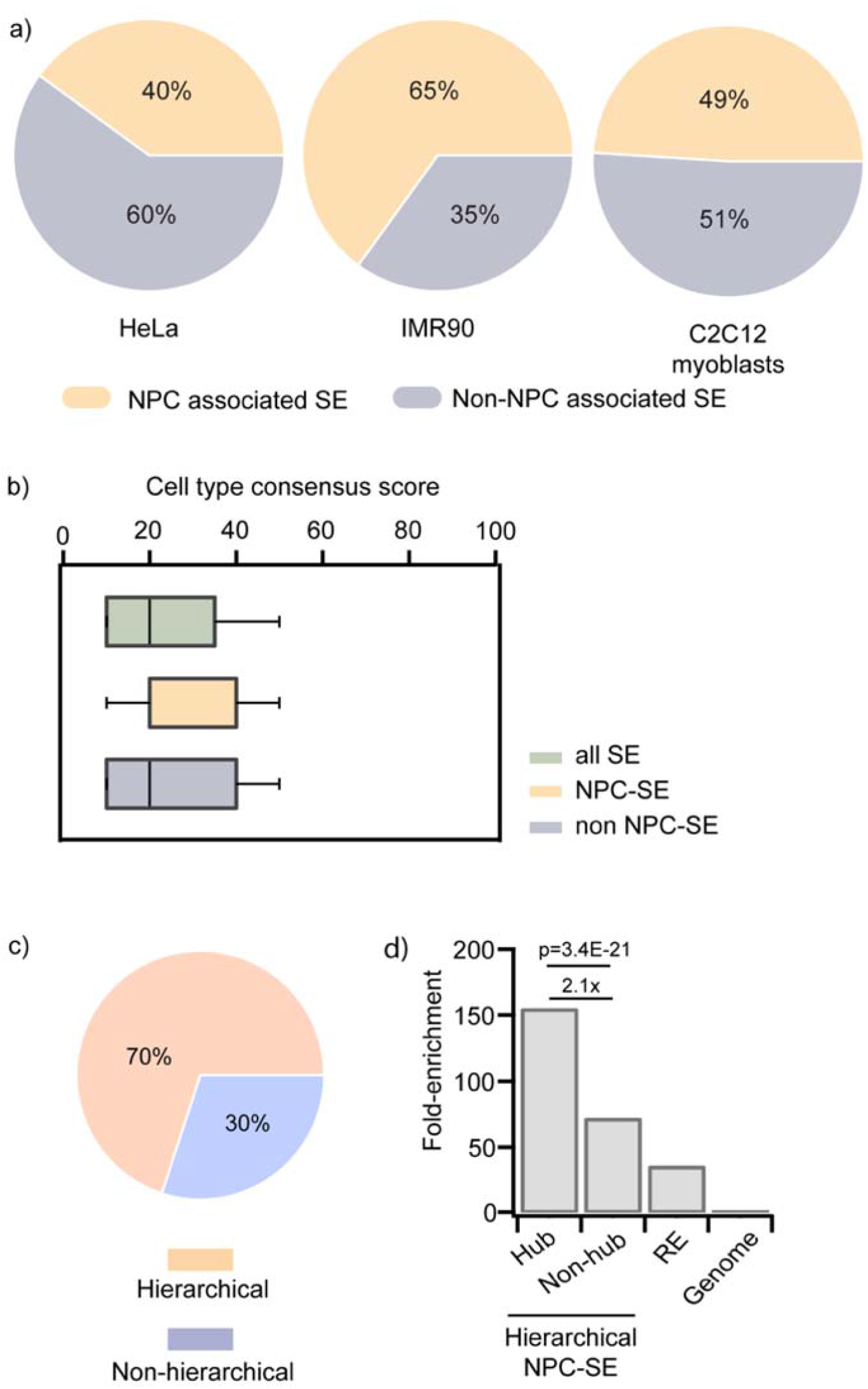
NPCs are consistently interacting with SEs across multiple cell types. a) Pie charts show the overlap of NPC-DamID peaks with SEs and non-SE regions for HeLa, IMR90, and C2C12 myotubes. b) Bar plots for cell type consensus score of all SEs, NPC-associated SEs (NPC-SE), and SEs not associated with NPCs (non-NPC-SE) in IMR90 cells. c) Pie chart showing that 70% of NPC-SE have hierarchical organization while 30% are non-hierarchical. d) The occupancy of NPC in four enhancer groups: hub (n=1,050) and non-hub (n=7,874) enhancers within hierarchical NPC-SEs, regular enhancers in IMR90, and randomly selected genome regions as control. *P* values were calculated using Fisher’s exact test.

A recent study revealed that a subset of SEs is hierarchically organized into hub and non-hub enhancers. The hub enhancers are enriched with CTCF and cohesin binding and play an important role in SE functional and structural organization ^20^. Based on this knowledge, we analyzed the structural organization of NPC-SEs in IMR90 cells. Intriguingly, a significant fraction of NPC-SEs (∼70.0%) have a hierarchical organization (Fig. 4c). Within hierarchical NPC-SEs, we observe that NPCs preferentially bind to hub enhancers compared to non-hub enhancers (2.1-fold, *P* = 3.4E-21, Fisher’s exact test, Fig. 4d). Interestingly, we further observe that the co-occupancy of NPC and structural proteins like CTCF, BRD4, P300, and PolII is markedly elevated at hub enhancers compared to non-hub enhancers (5.1-fold, 2.9-fold, 2.2-fold, and 2.2-fold; *P = 3.4E-08, P = 3.8E-21, P = 4.8E-13, and P = 2.4E-11*, Fisher’s exact test, Supplementary Fig. 4c-f). This result suggests that NPCs might have functionally related roles on hub enhancer elements in mediating the structural and functional organization of SEs through the recruitment of structural protein complexes (Supplementary Fig. 4g).

These results suggest that NPC-SEs are important for genome organization and folding, possibly through interaction or regulation of chromatin architectural proteins.

### NPCs are hubs for chromatin structural proteins

Next, we sought to understand the mechanism through which NPCs engage and regulate the activity of SEs. SEs have high occupancy of chromatin structural proteins like CTCF, mediator proteins, coactivators like P300, BRD4, cohesins, and PolII ^28^. Our finding that NPCs are associated with hierarchical hub enhancers (Fig. 4c-d) encouraged us to propose a model where NPCs interact with chromatin structural proteins and participate in maintaining the structure of SE domains at the nuclear envelope (Supplementary Fig. 4g). To test this possibility, we used proximity ligation assays (PLA) to assess the interaction between the NPC component TPR and structural proteins. PLA allows the identification of in-situ protein interactions (at a distance <40 nm) with high specificity and sensitivity. Several SE-associated proteins, including CTCF, PolII, Med1, BRD4, p300, and SMC3, show enriched PLA signals at the nuclear rim, indicating their interaction with TPR at NPCs (Fig. 5a and Supplementary figure 5a). Moreover, using TPR depletion as a negative control in PLA experiments (Fig. 5a), we show that PLA intensity (Fig. 5b) and the number of PLA puncta (Fig. 5c) significantly decrease in TPR knockdown cells, suggesting a true interaction between TPR and chromatin structural proteins.

**Figure 5:**
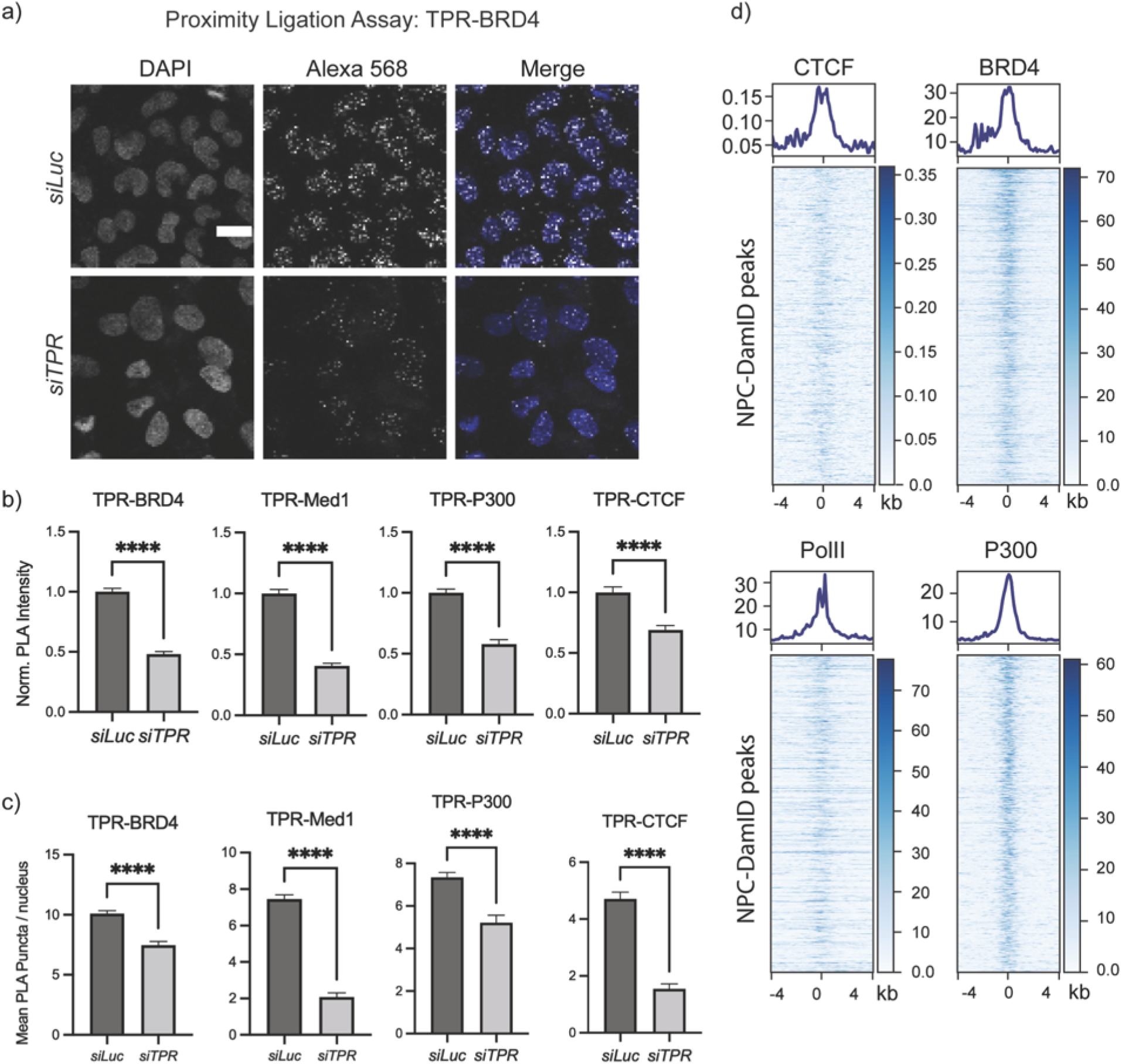
NPCs are hubs for chromatin structural proteins. a) Representative images of proximity ligation assay (PLA) between the nucleoporin TPR and BRD4 upon treatment with siRNA against Luciferase control (top panel) or TPR (bottom panel) in U2OS cells. Scale bar is 20 μm. Images are maximum intensity projections of confocal z-slices. b) Bar plot of mean fluorescence intensity of PLA signal per nucleus for TPR interaction with BRD4, Med1, P300, and CTCF in control and TPR knockdown cells. Data normalized to the average value of the control (siLuc) sample. c) Bar plot of the mean number of PLA puncta per nucleus for TPR interaction with BRD4, Med1, P300, and CTCF in control and TPR knockdown cells. For b and c, n>250 nuclei per condition, error bars represent a 95% Confidence Interval, and a t-test was used to calculate statistical significance (**** p< 0.0001). d) Distribution of reads of CTCF, BRD4, PolII, and P300 ChIP-seq centered around NPC-DamID peaks in IMR90 cells shown as a metagene profile and heatmap.

Further, we confirmed the overlap of NPC-associated chromatin with the DNA occupied by the structural proteins. To do this, we analyzed the distribution of the ChIP-seq reads for the structural proteins CTCF, BRD4, P300, and PolII compared to our NPC-DamID reads. We see a good correlation in the distribution of most of these proteins across the entire genome (Supplementary Fig. 5b). Further, looking at the distribution of the abovementioned structural proteins around the center of NPC-DamID peaks, we see specific enrichment for these proteins at NPC-linked chromatin regions (Fig. 5d). Together, the data suggest that NPCs are structural hubs for SEs, which may be maintained through interaction with chromatin structural proteins.

### Nups exhibit phase separation properties which aid their interaction with chromatin structural proteins

The observed association of SEs with NPCs raised the interesting question of how these interactions are established and maintained. Chromatin organization of SE regions is maintained through multivalent cooperative interactions between the intrinsically disordered regions (IDRs) of the mediator and transcription coactivator proteins ^21^. The disordered domains of proteins Med1, BRD4, and PolII, when present in high concentrations at SEs, can form multimeric intermolecular interactions leading to the formation of phase-separated condensates ^21^. NPCs are also highly enriched in completely or partially intrinsically disordered proteins. Like Med1, when Nup153, Nup98, Nup93, and TPR were tested through a disorder prediction computational tool called PONDR ^29^(Fig. 6a, Supplementary Fig. 6a, and data not shown), they showed large IDRs (45-85%).

**Figure 6:**
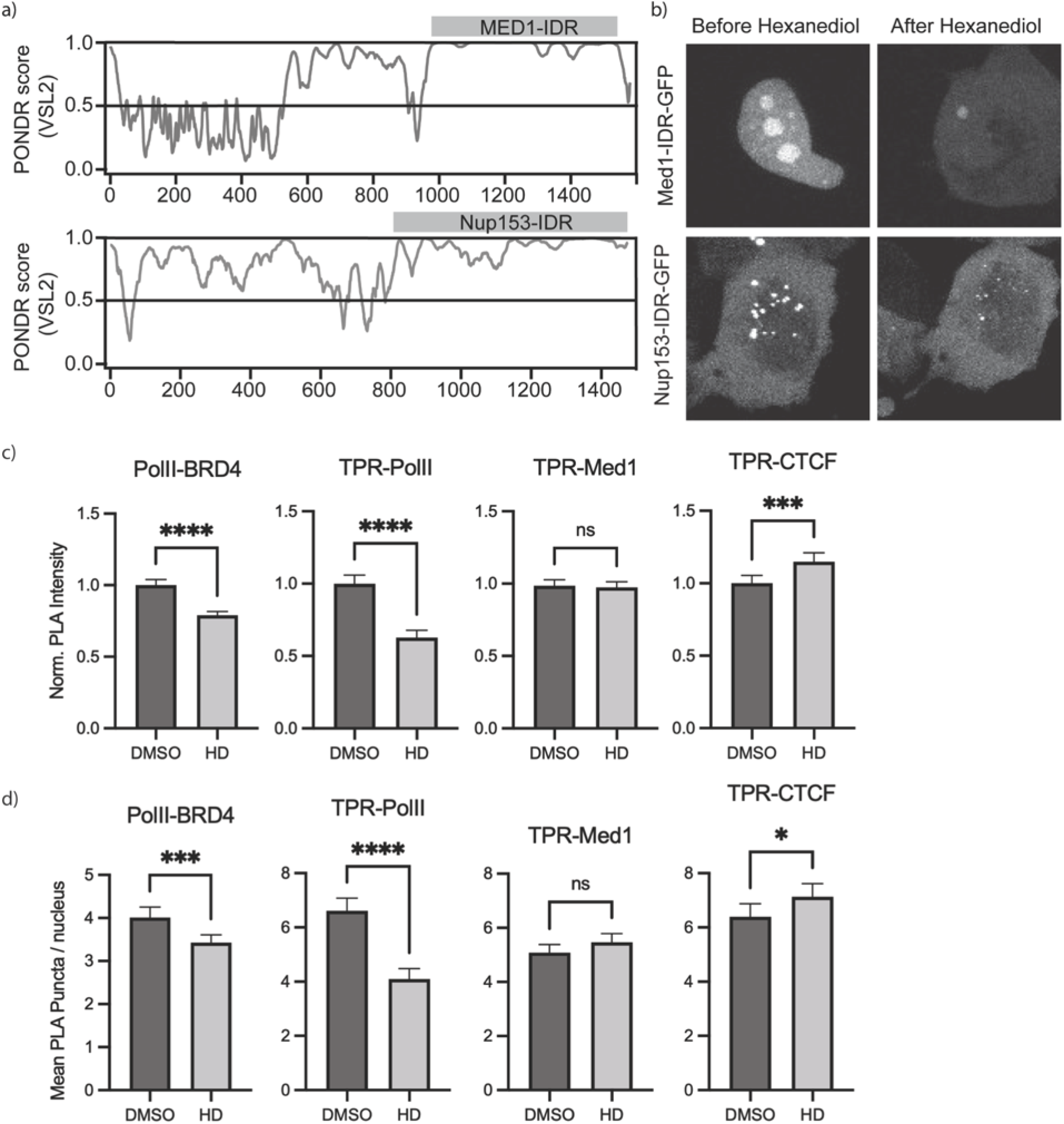
Phase separation properties of Nups are important for their interaction with other SE hub proteins. a) PONDR prediction for intrinsically disordered regions (IDR) in the structure of Med1 and Nup153. A score over 0.5 signifies a lack of structure. b) Confocal images of HeLa cells transfected with GFP fused with Med1-IDR or Nup153-IDR before and after 30s of treatment with 1.5% hexanediol (HD). c) Bar plots of mean fluorescence intensity of PLA signal per nucleus in DMSO and HD treated cells. The protein interaction pairs are mentioned on top of each plot. Data normalized to average PLA intensity of DMSO sample. d) Bar plots of the mean number of PLA puncta per nucleus in DMSO and HD treated cells for different protein interacting pairs mentioned on top of the plot. For c and d, n>300 nuclei per condition, error bars represent 95% Confidence internal, t-test was used to calculate statistical significance (ns not significant, * p<0.05, *** p<0.001, **** p< 0.0001).

These results encouraged us to test whether peripheral Nups in NPCs interact with chromatin structural proteins at SE condensate hubs through their IDRs (Supplementary Fig. 4g). The *in vitro* liquid droplet formation property of FG-containing Nups has been previously studied in the nuclear transport process ^30, 31^. However, its significance in chromatin organization has not been explored yet. We set out to compare the liquid droplet properties of Nup IDRs with Med1 IDR, which is involved in chromatin organization of SE regions through phase separation. We transfected the IDRs of Med1, Nup153 and Nup98 fused to GFP into cells and observed the formation of droplets. We saw that, similar to Med1, both Nup153 and Nup98 IDRs form droplets that disintegrated with hexanediol (HD), an alcohol commonly used to dissolve liquid-liquid phase-separated condensates (Fig. 6b and Supplementary Fig. 6b). The FRAP (fluorescence recovery after photobleaching) recovery time (τ) for Nup153 and Nup98 IDRs in a region of 1.5 µm^2^ was about 19 s compared to 11 s for Med1 IDR (Supplementary Fig. 6c). The diffusion coefficient for Med1 IDR was 0.13 µm^2^/s, as also determined by a previous study ^21^. The diffusion coefficients for Nup153 and Nup98 FG were found to be 0.076 and 0.077 µm^2^/s, roughly half of Med1 and similar to previously described components of liquid-like condensates ^21, 32^. From these results, we can conclude that Nups liquid-like droplets are sensitive to hexanediol disruption and are slightly less dynamic in diffusion than those of Med1.

To further test if the interaction between Nups and other SE proteins like BRD4 and PolII results in phase-separated condensates, we performed PLA experiments in the absence and presence of HD. We treated the cells with DMSO or HD for 20 min and probed for interactions through PLA. Upon quantifying the intensity of PLA spots and the number of PLA puncta, we found that TPR-PolII interaction decreased after HD treatment in a similar manner to the interaction between known co-phase separating condensate proteins BRD4 and PolII (Fig. 6c-d). However, PLA intensities and puncta number remained unchanged for TPR-Med1 and showed a slight increase for TPR-CTCF, indicating that not all interactions are phase separation dependent (Fig. 6c-d). Altogether, our results suggest the involvement of Nups in the organization of SE through interactions with chromatin structural proteins in phase-separated liquid droplets.

### Synthetic LacO arrays mediate the formation of Nup IDR interaction hubs that can recruit other chromatin structural proteins

Several studies have used *in vitro* purified IDR proteins at high concentrations to form hydrogel or liquid-liquid droplets to understand the biophysical and biochemical properties of interactions between IDRs ^21, 30^. The FG-repeats containing Nups, Nup98 and Nup153, have also been previously studied for *in vitro* hydrogel formation and its importance for selective permeability of nuclear transport receptors ^31^. There is a significant difference between *in vivo* physiological conditions and those used for *in vitro* phase separation experiments ^33^. Experimental conditions like temperature, concentration, pH, purity, and the microenvironment are detrimental to interactions between IDRs that form liquid droplets. Hence, we decided to use *in vivo* lac operator (LacO) arrays to probe the interaction between IDRs of Nups and structural proteins of SE hubs. U2OS cells containing the LacO array (harboring ∼50,000 LacO repeats) ^31, 32^ were transfected with fluorescently tagged IDRs fused to an NLS and LacI (mCherry-IDR-NLS-LacI), which led to the recruitment of a large number of molecules of mCherry-IDR-NLS-LacI at LacO sites (Fig. 7a-b). We observed concentrated *in vivo* interaction hubs inside the nucleus. We used this system with Nup153 IDR by fusing Nup153 IDR with mCherry, NLS, and LacI. Upon transfection of mCherry-N153-NLS-LacI (N153-LacI) to cells with the LacO array, we could observe a bright fluorescent spot with more intensity and larger size than mCherry-NLS-LacI (LacI), suggesting that the copy number of N153-LacI at LacO associated hubs is higher than LacI alone (Fig. 7b). A similar trend was observed for Med1-IDR-LacI spots. Additionally, LacI fused to N153 and Med1 could form multiple fluorescent regions throughout the nucleus compared to LacI alone, which leads to a single fluorescent spot. These results indicate likely cooperative and multivalent intermolecular interactions for N153-LacI and Med1-LacI inside the nucleus, critical for forming phase-separated droplets.

**Figure 7:**
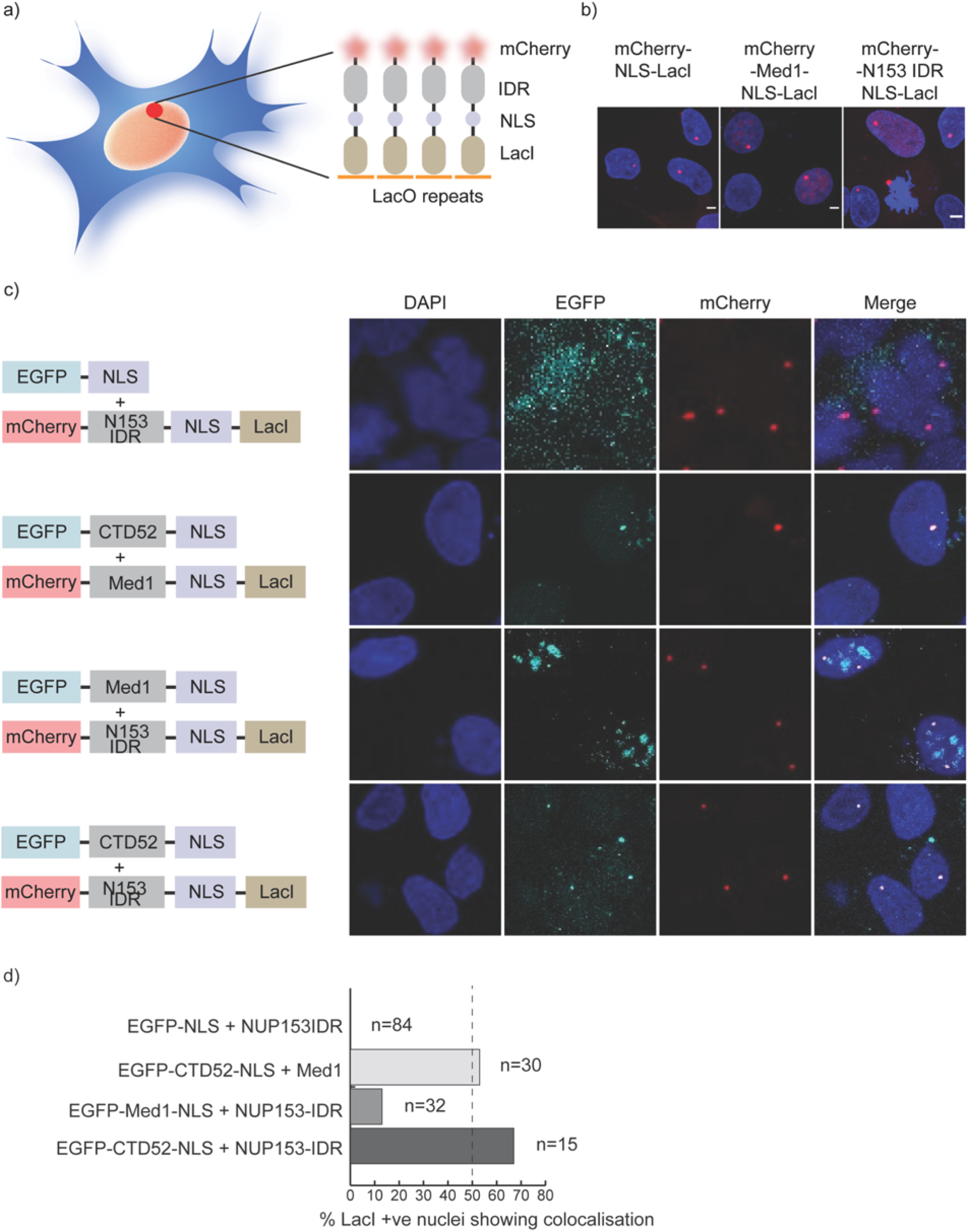
Synthetic LacO array mediated Nup IDR hubs can recruit other chromatin structural proteins. a) Scheme of a cell containing an array of LacO repeats (∼50,000) which can be decorated with the binding of LacI fused to a fluorescent protein (mCherry), IDRs, and nuclear localization signal (NLS). b) Confocal representative images of U2OS cells containing LacO arrays transfected with LacI fused to mCherry alone (mCherry-NLS-LacI), mCherry-Med1-IDR-NLS-LacI, or mCherry-Nup153-IDR-NLS-LacI. The scale bar is 1 µm. c) Confocal images of U2OS cells harboring LacO arrays and co-transfected with EGFP-NLS and mCherry-Nup153-IDR-LacI or EGFP-PolII CTD52-NLS and mCherry-Med1 IDR-NLS-LacI or EGFP-Med1-NLS and mCherry-Nup153-IDR-NLS-LacI or EGFP-PolII CTD52-NLS and mCherry-Nup153-IDR-NLS-LacI. d) Bar plot of the percentage of mCherry-LacI positive nuclei showing EGFP colocalization for images in c). The number of total nuclei counted is shown alongside the plot.

Next, we performed FRAP experiments to confirm the multivalent and cooperative interaction between LacI-IDRs at LacO hubs. We found that the dynamics of LacI alone spots were different than LacI fused to the IDR of either Nup153 or Med1. Upon fitting the FRAP curves with a reaction-dominant model ^33^, we found that Nup153 IDR and Med1 IDR fused to LacI had a reduction between 65-85% in the dissociation constant (k_2_) as compared to LacI alone (Supplementary Fig. 7). This suggests that the locally high concentration of Nup153 or Med1 IDR hubs at the LacO site are driven by IDR-IDR interactions via multivalent contacts, which would stabilize the association with cognate genomic DNA and other partner proteins. This result is relevant in the context of SE hubs which are formed through multiple homotypic and heterotypic interactions between IDRs of transcription factors and chromatin structural proteins ^21^.

After confirming homotypic multivalent interactions for the IDRs of Nup153 and Med1 fused to LacI, we investigated the potential of Nup153 IDR to interact with the known SE resident IDRs PolII and Med1. We again took advantage of the LacO-LacI system to form a hub for Nup153’s IDR to observe the recruitment of PolII and Med1. We co-expressed the CTD52 repeats of PolII or Med1 IDR fused to GFP in cells with mCherry-Nup153-IDR-LacI. We observed a colocalized fluorescent signal of GFP and mCherry for both PolIICTD52 (in 67% of cells) and Med1 IDR (in 13% of cells), while LacI alone cells showed only mCherry signal (no cells exhibited colocalization with GFP) (Fig. 7c-d). Similar colocalization was also observed when cells expressing mCherry-Med1-IDR-LacI were transfected with GFP-tagged PolII CTD52 (in 53% of cells). These results suggest that the Nup153 IDR domain, when present in high concentrations at the LacO site, can recruit the other partner proteins of SE hubs, PolII and Med1. This indicates that Nup153 functions similarly to Med1, previously shown to form phase-separated droplets at SEs sites to bind other structural proteins of SE. Altogether, these results support a model that peripheral Nups like Nup153 form phase-separated condensates at NPCs that compartmentalize and concentrate transcriptional coactivators and structural proteins at SE-regulated genes.

## DISCUSSION

This study presents a new method, NPC-DamID, to study NPC-mediated genomic interactions. NPC-DamID has several advantages over existing approaches to probe genome-wide interactions at the nuclear periphery. First, NPC-DamID eliminates any contribution of genomic interactions with Nups that are not at the NPCs. Previously it had been challenging to discern the difference between genomic interactions of Nups in the nucleoplasm and at NPCs due to the mobile nature of Nups used in other methods. Since NPC-DamID uses recombinantly expressed and purified Importinβ fused to Dam, specifically targeting NPCs in semi-permeabilized cells, it reports interactions exclusively at the nuclear periphery (Fig. 1a, d, e). We have shown through experimental and modeling data that NPC-DamID better reflects peripheral NPC-genome interactions than previously published datasets using Dam-Nup153 (Fig. 2 and Supplementary Fig. 2g-j). We have also demonstrated that the dominant mutant of Impβ (Dam-ΔN44 Impβ), which has a higher affinity to NPCs compared to wt Impβ ^25^, shows NPC-genome interactions similar to those of Dam-Impβ. In the future, NPC-DamID-seq can be expanded to map the NPC-associated proteome by fusing Impβ to APEX2 (a proximity labeling ascorbate peroxidase) ^34^, which will help discover novel interacting partners of NPCs involved in genome regulation. Second, our method allows us to probe NPC-genome interactions in any cell or tissue type since the technique does not rely on transgenic cell lines. To this end, we have mapped the interactions of NPC with genomic DNA in three cell lines and live human pancreatic islet tissue samples (Fig. 4a and Supplementary Fig. 3). In the future, NPC-DamID will be a valuable tool to understand better the influence of disease conditions like diabetes or cancer on genome architecture at the nuclear periphery. This is also possible since NPC-DamID can be done using as few as 50,000-200,000 cells. Third, NPC-DamID is not only independent of antibody and fixation as conventional DamID but also free from over-expression artifacts of Dam fused proteins. The concentration of Dam-Importinβ and the timing of the assay can be precisely controlled to avoid any artifacts. Fourth, the temporal resolution of NPC-DamID is higher than conventional DamID since there is no requirement of waiting 2-3 days to get enough expression of Dam fused protein. Through NPC-DamID, it is possible to obtain a snapshot of NPC-genomic DNA interactions at any point in time, for example, during the time course of cell differentiation (data not shown). Fifth, NPC-DamID sequencing can be combined with super-resolution imaging which is difficult with conventional DamID ^23^. In conventional DamID, Dpn7 (a DNA binding fragment of Dpn1) fused to a fluorescent protein is co-expressed with the Dam tagged protein of interest to allow imaging of the methylated DNA. This works well in cases like Dam-Lamin, when megabase pair long stretches of DNA methylation are expected. However, for sparse methylation patterns, as one would expect for NPC-associated DNA, the signal strength is too weak to perform high-resolution imaging, which requires high signal density. In NPC-DamID, we take advantage of the *in vitro* nature of the assay and use purified GFP or Flag-tagged Dpn8 labeled with a succinimide synthetic fluorescent dye, like Alexa 647 (Fig. 1e, f). This results in an increased number of dye molecules per Dpn8 molecule and gives us the flexibility of using synthetic dyes, which perform better in super-resolution methods like dSTORM. Thus, we can probe NPC association with chromatin through imaging at a single NPC resolution. A previously described model suggested that the heterogeneous composition of NPCs in a single cell could lead to differences in its interactions with chromatin ^35–37^. Our method, combined with imaging, will be useful in future studies to explore such models and understand the role of heterogeneity in NPC structure in genome organization. Furthermore, the *in vitro* nature of this method can allow biochemical manipulations to study the more specific nature of NPC-chromatin interactions. For example, in the future, we could map the distance-dependent interaction of the genome with NPCs by using Dam-Impβ with increasing linker lengths.

The association of Nups with the genome has been previously shown to be important for gene regulation. Recent evidence has indicated that the NPCs also mediate the structural organization of the genome^38–40^. However, a comprehensive understanding of the molecular mechanism underpinning such roles of Nups or NPCs remains unclear. Previous studies used specific Nup-targeted ChIP-seq or DamID-seq, thus having difficulty assessing whether the genomic associations shown represented binding to a mobile nucleoplasmic pool of Nups or NPCs at the periphery ^11, 13, 40^. These studies rely on DNA-FISH to test the localization of genomic interactions with Nups, which can only be tested for a few loci due to low throughput. With our new method NPC-DamID, we were able to probe genome-wide interactions of NPCs that happen specifically at the periphery in multiple cells and a tissue type to understand the common and unique nature of these interactions. We have found that the NPC-associated chromatin in various cell lines is primarily enriched in intergenic regions bound to H3K27Ac, followed by H3K4Me3 histone marks (Fig. 3a, Supplementary Fig. 3). This indicates that NPCs associate with regulatory enhancer and promoter regions, which falls in line with previous studies in human, metazoan, yeast, and mouse cells ^11, 14, 38, 40–42^.

Interestingly, in many recent studies in mice, drosophila, and yeast, Nup93 and Nup153 have also been shown to associate with repressed/facultative heterochromatin regions of genomic DNA ^13, 37, 43^. Our study found a higher association of actively expressed genes with NPCs, and only about 7% of NPC-DamID peaks overlap with Polycomb-associated repressed or quiescent genes (Fig. 3b). This result suggests that NPCs can associate with active and repressed regions, possibly through different Nups. It also incites the intriguing hypothesis that NPCs can vary in their composition and structure inside a nucleus, which leads to their association with different classes of genomic DNA and regulatory proteins. In the future, such models can be tested by combing NPC-DamID with super-resolution dual-color imaging to visualize the correlation between the structure and genomic associations of individual NPCs.

Another common genomic feature that associates with NPCs across multiple cells and tissue types is super-enhancers. This is consistent with previous work from our lab in which we have shown SE association with Nup153 and Nup93 through DamID-seq^11^. In the present study, we further explored the nature of NPC-associated super-enhancers (NPC-SE) compared to other SE regions.

Although super-enhancers are usually DNA regulatory elements that confer cell-type specificity to gene expression, a subset known as common super-enhancers is also active across multiple cell types. These common SEs are overrepresented in the pool of NPC interacting SEs identified in this study (Fig. 4b), and they have been implicated in establishing the early loop contacts during genome refolding in cell division ^19^.

Moreover, we found that NPC-SEs have hierarchical structural organization and preferentially enrich for hub enhancers (Fig. 4c-d). Hub enhancers are part of SEs that have a hierarchical organization. ^20^ This hierarchical structure is observed in about 25-60% of SEs in a cell. Hub enhancers are distinctly associated with CTCF, cohesin, and disease-associated genetic variants. Hub enhancers are also the primary elements responsible for SE functional and structural organization ^20^. In this way, they play an important role in dictating the chromatin landscape and regulating gene expression. These findings imply that the interaction of NPCs with SEs is important to the structural and functional organization of genomic DNA at the nuclear periphery through interactions with chromatin architectural proteins.

To explore the features and mechanisms for NPC-mediated genome organization, we tested the interaction of Nups to chromatin structural proteins, specifically those associated with SEs. We found through proximity ligation experiments that NPCs can interact with the SE proteins CTCF, Med1, Brd4, PolII, and cohesins. There is also significant enrichment of these proteins around the NPC-DamID peaks (Fig. 5). Together, these results strengthen the possibility of NPCs mediating the organization of SEs at the periphery through interactions with chromatin structural proteins. These results also corroborate recent studies showing the interaction of Nups like Nup153 or Nup93 with CTCF and cohesin ^40, 44, 45^.

We also explored how Nups at NPCs might interact with resident proteins of SE hubs. SEs are clusters of multiple enhancers which recruit transcription factors and coactivators containing intrinsically disordered regions (IDRs) in high concentration. The high concentration of IDRs allows multivalent interactions, leading to the formation of phase-separated condensates. This mechanism is important to ensure robust and high gene expression of cell identity genes. The peripheral Nups at NPCs, like Nup153, also have long IDRs, which we show in this study form phase-separated droplets *in vivo* (Fig. 6a-b). Nup153 is known to have at least 32 copies at individual NPCs ^35^, which likely facilitates high local concentration for phase separation. Since individual NPCs have been predicted to have different compositions within a single cell and are also known to vary in their composition from one cell type to another ^35^, SE-interacting NPCs may have even more than 32 copies to facilitate the interaction with SEs. In the future, high-resolution imaging could be done to test this hypothesis. We show that phase-separated Nup153 droplets can co-partition with IDRs of other SE resident proteins like Med1 and PolII (Fig. 7). This observation suggests that NPCs can be structurally and functionally part of SE hubs. At the same time, it raises multiple questions regarding the importance of NPCs in regulating this subset of SEs. Do NPCs act as a scaffold to organize SE hubs? How different are the organization and function of NPC-associated SEs compared to others? Why do certain SEs tend to associate with NPCs? In this regard, the gene-gating model, in which positioning a gene at NPCs facilitates the rapid transport of its mRNA to the cytoplasm ^46^, is more probable. Recent evidence from a study of an NPC-associated oncogenic super-enhancer for the MYC gene showed that SE positioning at the periphery facilitated the export of mRNA transcripts in colon cancer cells in response to WNT signaling ^17^. It will be interesting to determine if gene-gating holds for all NPC-associated SEs and non-cancer cells.

Interestingly, nuclear basket protein TPR is known to interact with the kinases ERK2 and CDK1 ^47, 48^, and Nup-SEs were shown to enrich for the motif for AP1-like transcription factors ^11^. It will be interesting to study if kinase interactions with TPR mediate the phosphorylation of AP1 transcription factors to facilitate gene expression of NPC-SEs.

In conclusion, NPC-DamID has allowed us better to understand the NPC’s role as a gene regulator. We have learned the common and unique features of NPC-interacting genomic regions across multiple cell types. Further, we have shown the role of disordered domains in Nups in the structural organization of SE hubs through phase separation. Moreover, further studies will be needed to better understand other interaction partners (protein or RNA) of NPCs that mediate the association of NPCs with the genome.

## Supporting information

Table S2: coordinates of NPC- and non-NPC SEs and transcription level of their associated genes.

Table S1: gene annotations around NPC-Dam peaks.

Table S3: coordinates of hub and non-hub NPC-SEs.

## ACKNOWLEDGEMENTS

This work was supported by M.H.’s NIH R01 grants (NS096786, GM126829) and Salk Cancer Center Support Grant P30 CA014195. M.H. also received financial support from the W.M. Keck Foundation and the NOMIS Foundation. Further, M.H. received support from the AHA-Allen Initiative in Brain Health and Cognitive Impairment award made jointly through the American Heart Association and The Paul G. Allen Frontiers Group (19PABH134610000).

S.T. and J.C. were supported by Salk’s Women & Science Awards. S.T. also received financial support from the Hewitt Foundation fellowship, and J.C. is a Paul F. Glenn Biology of Aging fellow. J.H. was supported by the National Natural Science Foundation of China (31871317 and 32070635).

We thank Roberta Schulte for assistance with in vitro transport assays, for comments that greatly improved the manuscript, and for helping refine the figures presented in this work. We thank Shefali Krishna for creating the diagram for the NPC-DamID method, for her input on super-resolution microscopy analysis, and her insightful comments on this manuscript. We thank all members of the Hetzer lab for helpful discussions of these research ideas and their thoughtful comments on this manuscript. We are also grateful to Salk’s core facilities for their assistance. Specifically, we thank the Next Generation Sequencing Core (NGS) for sequencing our DamID and RNA NGS libraries, the Advanced Biophotonics Core for assistance with super-resolution microscopy, and the Razavi Newman Integrative Genomics and Bioinformatics Core (IGC) for their input on analysis methods for DamID experiments.

## AUTHOR CONTRIBUTIONS

S.T: conceived, designed experiments, collected data, performed analysis, and wrote the paper. J.S.C.: collected data, performed analysis, addressed requested revisions, and wrote the paper revisions. J.X. designed, performed, and analyzed experiments related to nucleocytoplasmic transport assays and reviewed the paper. F.C. and J.H. analyzed the hierarchical organization of NPC-DamID peaks and reviewed the paper. R.S.: analyzed the data for PLA and LacO array experiments and reviewed the paper. M.W.H: conceived, supervised, and reviewed the paper.

## DATA AVAILABILITY

The Gene Expression Omnibus accession number for the data generated in this study is GSE176106. DamID-seq data for IMR90, Nup153 was used from GSE87831. LaminB1 DNA binding profile data was used from GSM1541019. CTCF ChIP-seq data from SRR639078 & SRR639079. PolII ChIP-seq data from GSM1055822. BRD4 ChIP-seq data from SRR2748697 and p300 ChIP-seq data from GSM1055812. DamID-seq data for Nup98 was used from GSE83692.

## Materials & Methods

### Recombinant expression and purification of the proteins

The plasmids for nuclear transport factors (Impβ, Impα, NTF2, Ran, NLS-MBP-GFP cargo) were received from Prof. Edward Lemke’s laboratory. The plasmid contained intein-his tags for Ni-NTA column purification. Later, the tag was removed by incubating protein in 100 mM beta-mercaptoethanol for 12-14 h at RT in the presence of Complete protease inhibitor cocktail. The details of plasmids are also listed here ^49^. The purification steps for the proteins were followed according to Milles et al. ^50^ Dam was cloned in pet28 vector and expressed for 10 h at 18°C in HMS174(DE3) E.Coli cells. Dam-Impβ and Dam-ΔN44Impβ were cloned into pTXB3 plasmid with intein-His tag. It was also expressed in HMS174(DE3) cells for 10 h at 18°C. Ni-NTA purification was done as per manufacturer’s guideline. After purification through Ni-NTA column, intein-His tag was removed through beta-mercaptoethanol and second Ni-NTA column purification was done. The protein was then concentrated and purified through size exclusion column chromatography.

For loading GDP to Ran wwe incubated Ran with 50 X molar ratio with respect to protein with GDP in 20 mM HEPES (pH 8.0), 200 mM NaCl, 2 mM DTT and 5 mM MgCl_2_ for 1 h at 4 C. After that we quenched the reaction with 25 mM MgCl_2_ and overnight dialysis was against 20 mM HEPES, 200 mM NaCl, 2 mM DTT and 5 mM MgCl_2,_ 2 mM GDP.

We received ER295 and ER295(DE3) E. Coli strain and pet28-DpnI winged helix domain (Dpn8, residues 183-254 aa) from Prof. Michael Bochtler. We cloned flag tag and also GFP tag into the plasmid Pet28-Dpn8. The protein was expressed in ER2925 DE3 E. Coli cells and purified following the step from Mierzejewska et. al. ^22^

### *In vitro* lambda DNA protection assay (enzyme assay for Dam proteins)

Unmethylated lambda DNA was used for protection assay to test the activity of Dam and Dam fused proteins. 1 μg of LDNA was incubated with different concentration of Dam and Dam fused protein along with 7.5 µM SAM (s-adenosyl methionine) in transport buffer (20 mM HEPES, 110 mM KOAc, 5 mM NaOAc, 2 mM MgOAc, 2 mM DTT, pH 7.3) for 30 min at RT. The enzyme was then deactivated by heating samples at 65°C for 10 min. Then 1X DpnII NEB buffer was added along with 1 μl of DpnII. The samples were incubated at 37°C for 30 min.

The samples were heat inactivated and mixed with agarose gel loading dye. The samples were loaded on 0.8% agarose gel with EtBr to assess the activity of the proteins.

### Protein labeling through succinimide ester dyes

We labelled wt Impβ, Dam-Impβ and Dpn8 with succinimide Alexa dyes with free -NH2 group in the proteins. The dye stock solution was made in anhydrous DMF solution and stored in lypophillized form at -20°C. The labeling was done is 5X molar excess of dye in sodium phosphate buffer, 2 mM DTT, pH 7.5 for 2h at RT. Later the reaction was dialyzed or purified through size exclusion column to remove excess dye. The success of reaction was checked by run samples on SDS-PAGE and scanning the gel on fluorescent gel scanner.

### Nuclear import assay

The buffers used were PBS, permeabilization buffer (50 mM HEPES, 50 mM KOAc, 8 mM MgCl_2_, pH 7.3), and transport buffer (20 mM HEPES, 110 mM KOAc, 5 mM NaOAc, 2 mM MgOAc, 2 mM DTT, pH 7.3). The HeLa cells were washed for 3 × 2 min with PBS, followed by a 2-min wash with permeabilization buffer, followed by a 5-min permeabilization with digitonin at a concentration of 50 μg/ml. The digitonin was subsequently removed by washing for 3 × 3 min with transport buffer. After the final wash, HeLa cells were treated with an active nuclear import reaction mix containing a fluorescent import cargo probe (NLS-MBP-GFP), importin-β (1 µM), Importinα (1 μM) RanGDP (5 μM), NTF2 (4 µM), and an energy regenerating system (2 mM GTP, 0.1 mM ATP, 4 mM creatine phosphate, and 20 U/ml creatine kinase) in transport buffer. Import reactions proceeded at room temperature for 20 min before the cells were fixed with a 4% PFA solution for 15 min and washed 3 × 2 min with PBS. Cells were then imaged using a Zeiss LSM 700 confocal laser scanning microscope.

### NPC-DamID assay

For NPC-DamID assay, cells were grown in 6 welll plate with 70-80% confluency. The media was washed 3X with 1X PBS at RT. After washing, cells were incubated with permeailization buffer for 3 mins followed by digitonin permeabilization for 5 min at RT. Digitonin concentration ranged from 20-30 μg/ml for semi-permeabilization. The concentration was optimized for different cells through checking immunofluorescence for antibody against lamin in semi-permeabilized cells. The digitonin was then washed by 3X wash with transport buffer for 3 min each. Then transport buffer wit cocktail of proteins and SAM required for the assay was added, 5 units of Dam or Dam-fused proteins, 4 µM NTF2, Importinα (1 μM), 7.5 μM SAM. For assays with RanGDP/GTP, RanGDP (5 μM), and an energy regenerating system (2 mM GTP, 0.1 mM ATP, 4 mM creatine phosphate, and 20 µ/ml creatine kinase) was also added. The cells were incubated with transport mix for 20 min at RT with occasional gentle rocking to mix the reagents. After 20 min, the cells were washed with 1X PBS for 3X, 3 min each.

### NPC-DamID imaging through Dpn8

For super-resolution and confocal imaging cells were grown on glass coverslips or ibidi glass bottom chambers (for dSTORM imaging) with #1.5 thickness. After performing NPC-DamID assay, the reagents of the assay were thoroughly washed with 1X PBS at RT. Then cells were fixed with 2.4% PFA for 10 min and cells were permeabilized with 0.5% triton for 3 min. The cells were then blocked with 5% BSA for 1 h and incubated with 2 µM of GFP-Dpn8/Flag-Dpn8 solution in 5% BSA for 30 min at RT. The cells were washed 3 times with 1 X PBS and costained with DAPI or Nup153 antibody for NPC marker. The cells were then post-fixed with 2.4% PFA for 5 min and mounted in vectasheild for confocal imaging. For super-resolution imaging, cells were exchanged in fresh STORM photo switching buffer (1X PBS, 100 μM MEA cysteamine hydrochloride), 10% glucose, 5 µl catalase, 0.6 mg/ml glucose oxidase, pH 8). dSTORM single color and dual color was performed on ONI nanoimager (oxford nanoimaging instrument) with 640 nm and 488 nm excitation laser and 100X 1.49 NA Olympus objective. Up to 30,000 frames were collected with frame rate of 50-75 Hz.

### Cell culture

HeLa and U2OS cells were cultured in DMEM media supplemented with 10% FBS. IMR90 cells were cultured in 15% FBS, 1X NEAA (non-essential amino acids) in DMEM media. C2C12 cells were cultured in DMEM media supplemented with 20% FBS. To convert C2C12 myoblast to myotubes, C2C12 myoblasts were allowed to become confluent and then DMEM medium was supplemented with 2% horse serum instead of FBS. Myotubes were collected for the assay at 14 days. Pancreatic islets were bought from Prodo laboratories Inc. Islets were maintained in cell culture at 37 °C, 5% CO_2_ in PIM(S) Complete media for up to 7 days. Islets for NPC-DamID assay were disrupted into single-cells using TrypLE just before the assay. The assay was performed in suspension in tubes.

### NPC-DamID sequencing

Cells after NPC-DamID assay were scraped from 6 well plate and used for genomic DNA isolation. Qiagen gDNA isolation kit was used using manufacturer’s protocol. DNA was eluted in 500 µl of water or elution buffer. The DNA was then concentrated using rotatory evaporator if eluted in water otherwise precipitated with ethanol to dissolve in 50 μl of water so that concentration is about 500 ng/µl. The DNA was then prepared for library preparation following the published protocol for DamID-seq^51^. The library for DNA sequencing was prepared using Roche Kapa Hyper-prep DNA kit following manufacturer’s protocol. The quality of the DNA libraries were checked using Tapestation bioanalyzer and Quibit. The libraries were usually sequenced with single-end 50 bp sequencing.

Libraries were pair-end sequenced on illumina Nextseq (75 or 150 cycles) and basecalls performed using CASAVA version 1.8.2. Fastq sequence alignment to the reference genome (hg19 for human samples and mm10 for mouse) was done using STAR (v02.02.01) for both DamID-seq and RNA-seq datasets.

DamID peaks were found by merging replicates and using the findPeaks program in HOMER (add ref PMID: 20513432) with the following options “-style histone -o auto -size 2500 -minDist 2500 -tbp 0 -inputtbp 0 -fragLength 1 -inputFragLength 1 -F 2 -C 0 -ntagThreshold 100” using the Dam alone DamID experiments as background. DamID peaks identified were annotated using HOMER’s annotatePeaks.pl function with reference genome hg19, NPC-associated genes were defined as those whose gene body or promoter regions overlapped with NPC-DamID peaks.

For RNA-seq experiments, HOMER was used to create tag directories and annotation. Reads Per Kilobase of exon per million mapped reads were calculated using HOMER across annotated gene exons (NCBI RefSeq annotation).

For filtering, all reads which uniquely map and had MAPQ>10 were selected for downstream analysis. Duplicates were removed using PICARD tools. The peaks overlapping blacklisted region of genome were also removed.

HOMER (v4.11) was used to process alignment files to generate normalized bedGraphs and bigwigs for the UCSC Genome Browser. For other coverage measurements, DeepTools, v3.1.3 (BamCoverage) was used.

### Identification of Super-enhancers from H3K27Ac ChIP-seq

The identification of super-enhancers was done using H3K27Ac ChIP-seq data. SE identification from these was performed by HOMER’s findPeaks function using “-style super” as described in http://homer.ucsd.edu/homer/ngs/peaks.html#Finding_Super_Enhancers.

Briefly, peaks are identified and those within a given distance are combined into larger regions (12.5 kb). The super enhancer signal of each of these regions is determined by the total normalized number of reads (input subtracted). These regions are sorted, normalized to the highest score and the number of putative enhancer regions, and super enhancers are identified as regions past the point where the slope is greater than 1.

### Super enhancer consensus scoring

For super-enhancer consensus scoring, we downloaded the annotated SE sequences for 10 different cell lines from super enhancer database (SEA) ^52^. Then all SE sequences were merged and used as total number of SEs. Then each of them was tested for co-occurrence in 10 cell types and scored (s) using formula 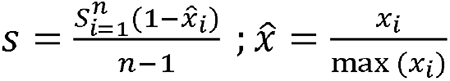 where x indicates cell/tissue type specificity and n denotes the number cell/tissue types ^19^.

### Super enhancer hierarchy determination

Super-enhancer hierarchical score determination using NPC-DamID peaks and Hi-C already published data for IMR90 was done using the methodology described previously by Huang et.al. ^20^ For correlation with the RNA expression, we used RNA seq data for IMR90 and HeLa cells and selected the closest gene in distance from the super-enhancer as the related gene for calculating the expression.

### PLA experiment

We performed PLA experiments using Duolink PLA kit from Sigma-Aldrich. We followed the protocol from the manufacturer for the experiments. For imaging we used Leica SP8 confocal microscope. For quantitation, DAPI signal from maximum projected images were used to create nuclear masks in Fiji (ImageJ). PLA signal (Integrated density) was measured within the nuclear mask for individual nuclei. To calculate normalized PLA intensity, all values for a given pair of PLA interactors were normalized to average PLA intensity of control group (siLuc or DMSO). For counting number of PLA Puncta, we used the ‘Find Maxima’ function on Alexa565 channel of maximum projected images and adjusted threshold to restrict maxima identification within nuclear masks. Threshold was kept constant between control and treatment groups for a given pair of PLA interactors allowing us to compare relative differences. All graphs and statistics were generated using Graphpad Prism.

### FRAP experiment

FRAP experiments were performed on Leica SP8 using the module for FRAP. To measure the FRAP dynamics of GFP fused to Nup-IDRs and Med1-IDR we acquired 90 frames every 1.3 s with 5 frames before bleaching. For m-Cherry labeled LCD-LacI or LacI, we acquired 200 frames every 2 s and 5 frames before bleaching. The average of first 5 frames before bleaching was used to calculate the background fluorescence for the baseline. We chose a circular spot of radius 1.5-2 µm for photobleaching.

We followed the steps of analysis from Chong et. al. ^33^We corrected xy-drift of bleached spot using ImageJ plugin “Template matching and slice alignment”. We normalized the mean intensity of bleach spot and whole nucleus at time t to pre-bleaching baseline intensity. After this normalization we normalized to relative bleach spot intensity to nuclear intensity. Then we calculated the difference between normalized FRAP intensity before and after photobleaching. Finally, we normalized this difference in intensity to 100% to plot the curves. The reaction-dominated fitting to LacI curves was also done as previously described^33^.

## SUPPLEMENTARY FIGURES

**Supplementary Figure 1.**
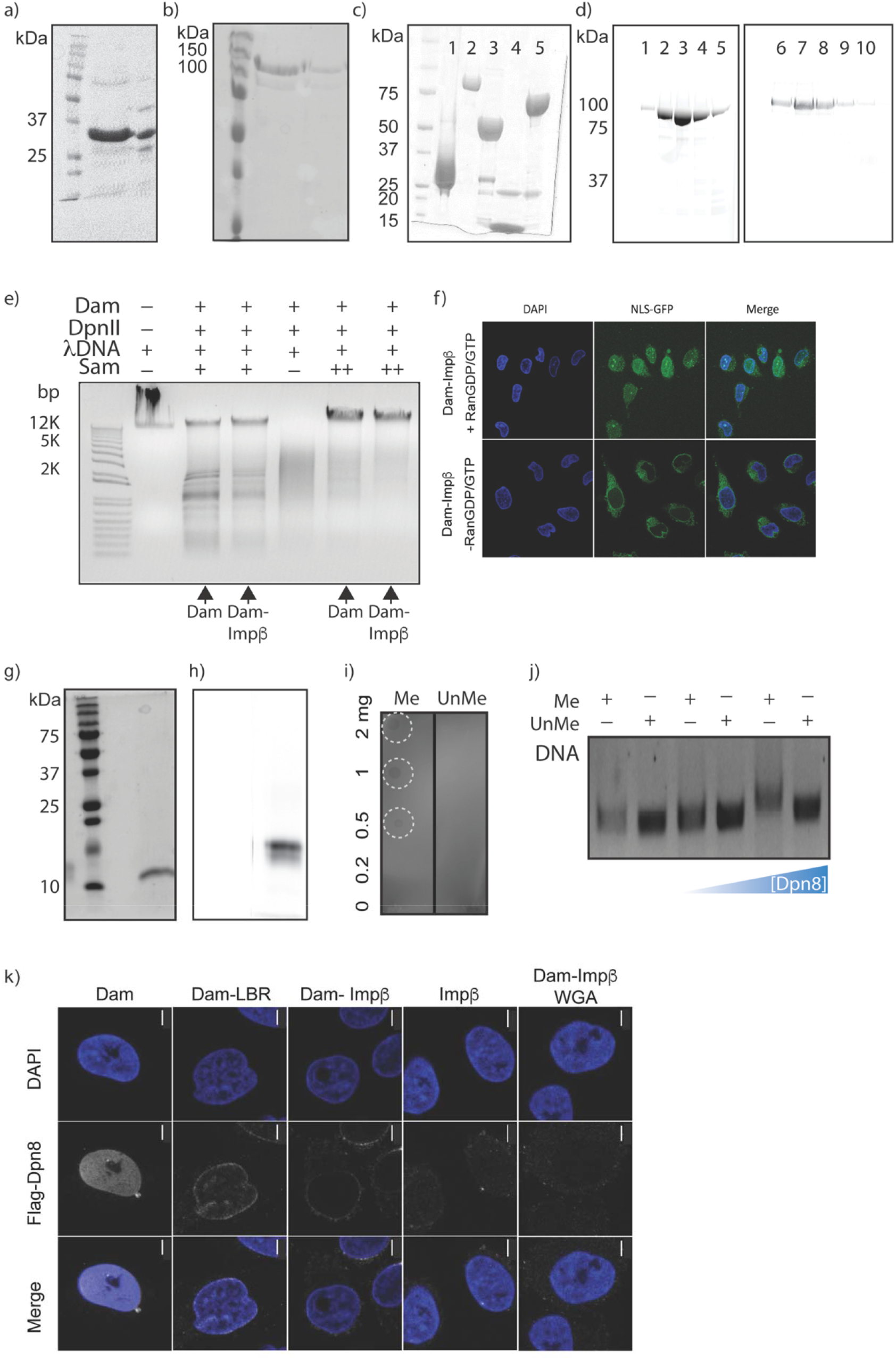
a) Coomassie gel for recombinantly expressed and purified Dam protein. b) Coomassie gel for recombinantly expressed and purified Dam-Impβ protein. c) Coomassie gel for recombinantly expressed and purified nuclear transport factors-Ran (Lane 1), Impβ (Lane 2), Impα (Lane 3), NTF2 (Lane 4), NLS-GFP-MBP cargo (Lane 5). d) Fluorescent gel scan of elutions for labeled Impβ (Lanes 1-5) and Dam-Impβ (Lanes 6-10) with Alexa 488 succinimide dyes after size exclusion chromatography e) Activity assay for Dam and Dam-Impβ with varying concentrations of SAM-5 μM (+) and 7.5 μM (++). f) Confocal images of the nuclear transport assay of NLS-MBP-GFP using Dam-Impβ in the presence and absence of RanGDP/GTP. The scale bar is 15 µm. g) Coomassie gel for recombinantly expressed and purified Dpn8. h) Fluorescent gel scan for Dpn8 labeled with Alexa 488. i) Dot blot assay for binding specificity of GFP-Dpn8 with methylated (Me) and unmethylated (UnMe) lambda DNA. j) Gel shift assay for Dpn8 binding to methylated lambda dsDNA (Me DNA) with increasing concentration of Dpn8. k) Confocal images of Hela cells labeled with Flag-Dpn8 (2μM) after NPC-DamID assay. Dam alone shows a uniform nucleoplasmic signal while Dam-LBR and Dam-Impβ show enrichment of signal at the nuclear periphery for LADs and NPC-interacting DNA, respectively. Impβ alone and Dam-Impβ with WGA (blocking agent for NPCs) show no Dpn8 signal. The scale bar is 5 µm.

**Supplementary Figure 2.**
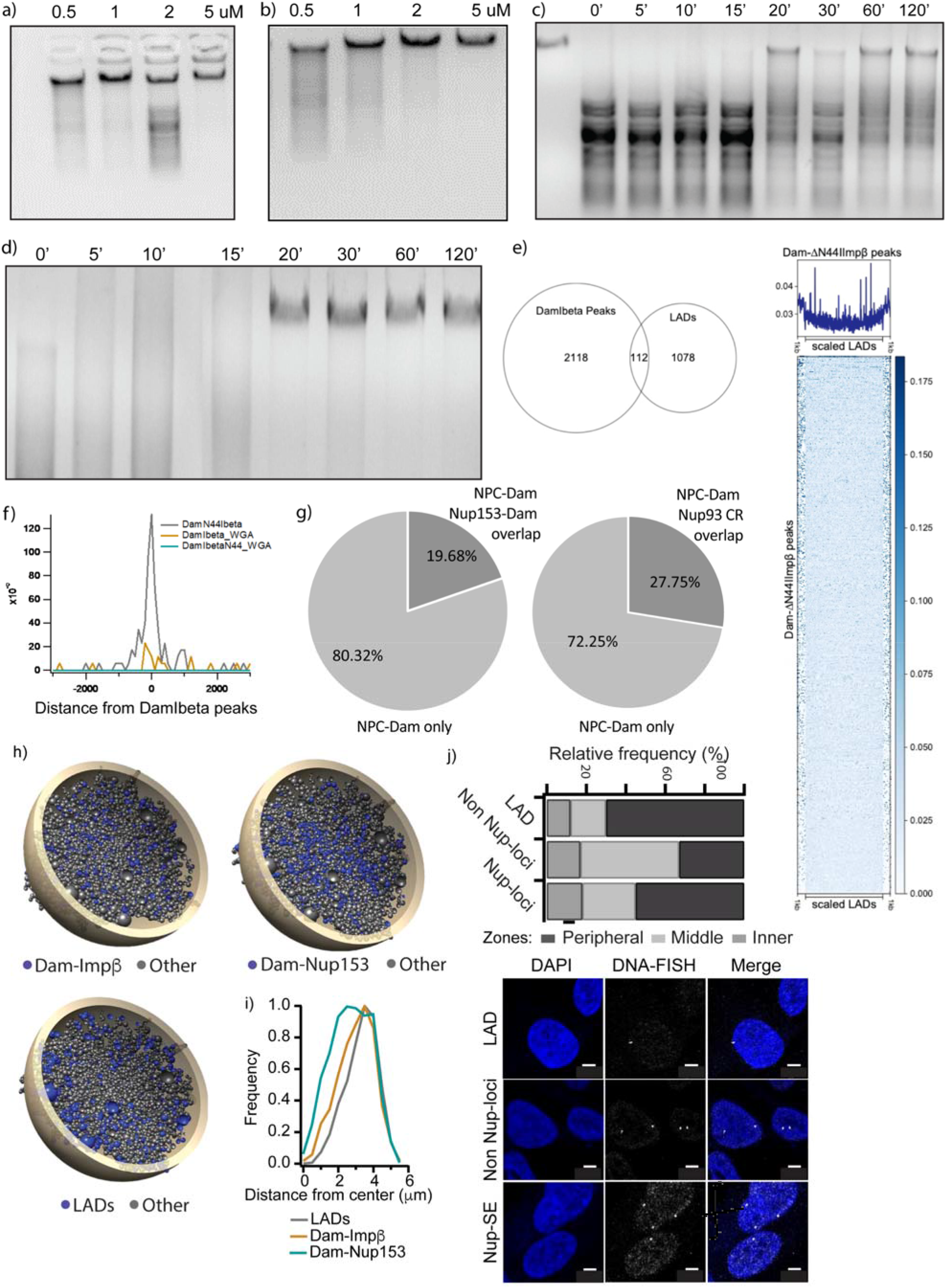
a) EtBr-stained agarose gel for testing enzymatic activity of Dam. The unmethylated lambda DNA (1 µg) was incubated with an increasing concentration of Dam in the presence of SAM for 30 min at RT. After 30 min, DpnII was added to digest the remaining unmethylated DNA after the enzyme reaction. We observe the efficient protection of lambda DNA from digestion by DpnII through methylation by Dam. b) Same as a) for Dam-Impβ. c) Time-dependent activity of Dam (reaction set up was same as in a). Dam (5 units) requires at least 20 min at RT for complete methylation of 2 µg of lambda DNA. d) Same as c) for Dam-Impβ. e) Left, Venn diagram of the overlap between NPC-DamID peaks and LADs in IMR90. Right, heatmap and metagene profile showing the distribution of Dam-ΔN44Impβ around LAD regions. f) the distance-dependent distribution of the read density of Dam-ΔN44Impβ, Dam-Impβ+WGA, Dam-ΔN44Impβ+WGA centered around Dam-Impβ peaks. g) Proportion of NPC-DamID peaks overlapping with Nup153-DamID peaks or Nup93 cut&run peaks in IMR90 cells. h) Snapshot of the 3D computational model of IMR90 using Chrom3D with LADs as a peripheral nuclear constraint. Dam-Impβ peaks and Dam-Nup153 peaks were then overlapped to observe their spatial distribution. i) Frequency histogram for the distance of the beads corresponding to genomic loci of the LADs, Dam-Impβ, or Dam-Nup153 peaks from the center of the nuclei computed through Chrom3D models. j) DNA-FISH-based determination of the nuclear localization of loci shown to interact with LADs, the NPC, or neither.

**Supplementary figure 3:**
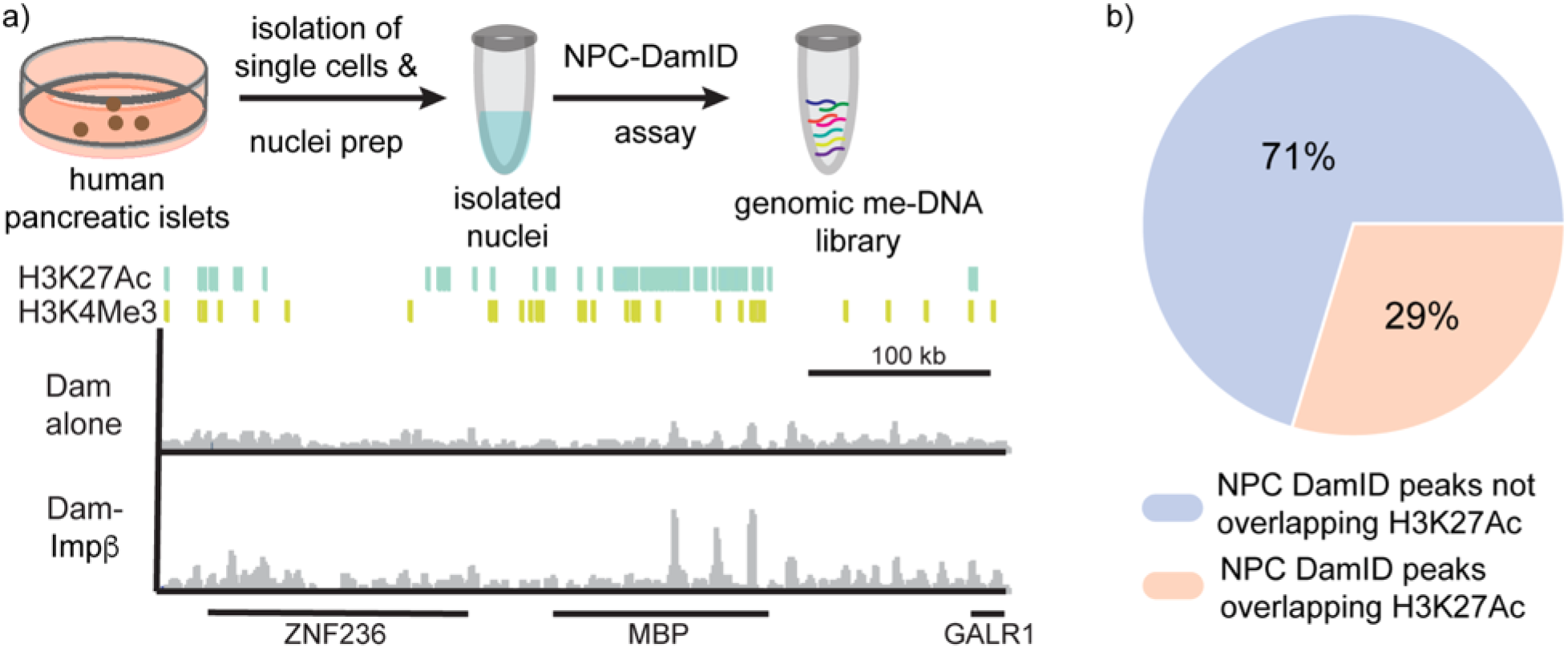
NPC-DamID is also applicable to tissue samples. a) Top, schematic representation of the NPC-DamID assay for live human pancreatic islet tissues. Bottom, the normalized profile for Dam alone and Dam-Impβ along with the tracks for histone marks-H3K27Ac and H3K4Me3 for human pancreatic islets are shown. b) Pie chart for H3K27Ac overlap with NPC-DamID peaks for human pancreatic islets.

**Supplementary Figure 4:**
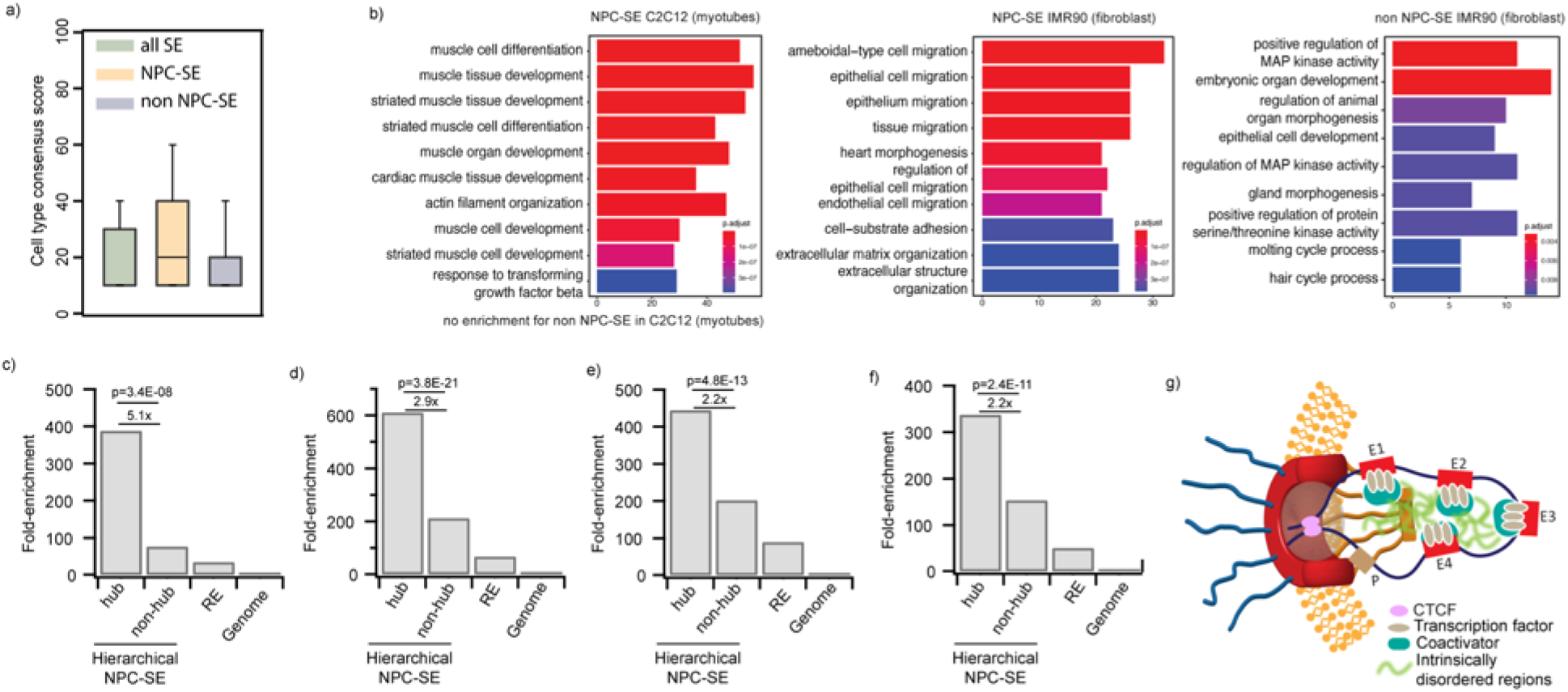
a) Bar plots for cell type consensus score of all SEs, NPC-associated SEs (NPC-SE), and SEs not associated with NPCs (non-NPC-SE) in HeLa cells. b) GO biological process enrichment analysis for genes associated with NPC SEs and non-NPC SEs in IMR90 (fibroblasts) and C2C12 (myotube) cells. c-f) Bar plots showing fold-enrichment for CTCF (c), BRD4 (d), P300 (e), PolII (f) co-occupied NPC-associated hierarchical SEs (NPC-SE) with hub, non-hub, and regional enhancer (RE) over the rest of the genome. g) Scheme for the model of NPCs acting as structural hubs for the organization of SEs through interactions mediated by disordered regions of Nups, transcription factors, and coactivators.

**Supplementary Figure 5.**
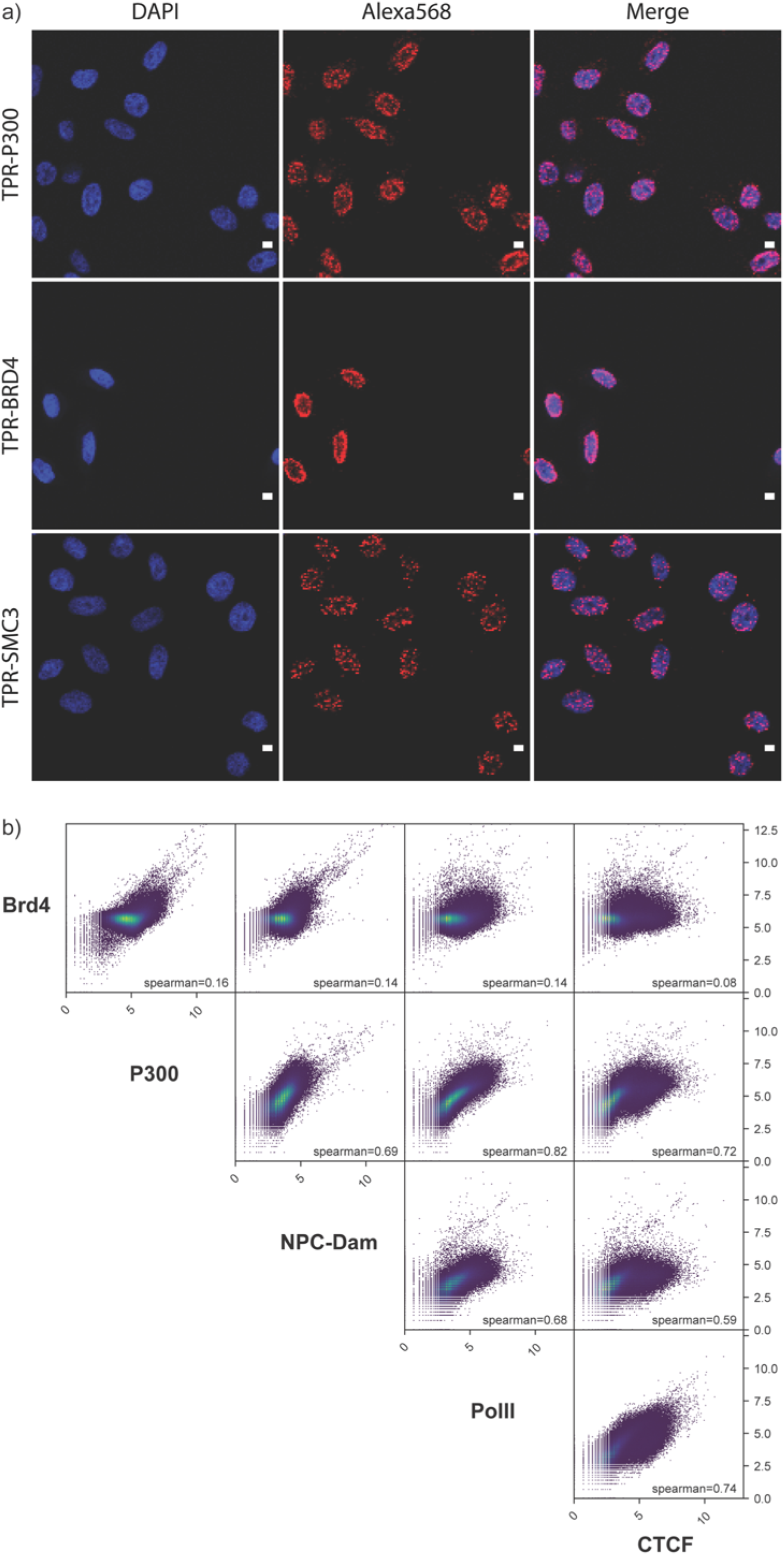
a) Confocal images for PLA experiment between TPR and chromatin structural proteins P300, BRD4, SMC3. The scale bar is 10 μm. b) Scatter plots comparing co-occupancy between CTCF, BRD4, POL II, P300, and NPC-DamID across the genome. The Spearman correlation coefficient is also shown along with each comparison.

**Supplementary Figure 6.**
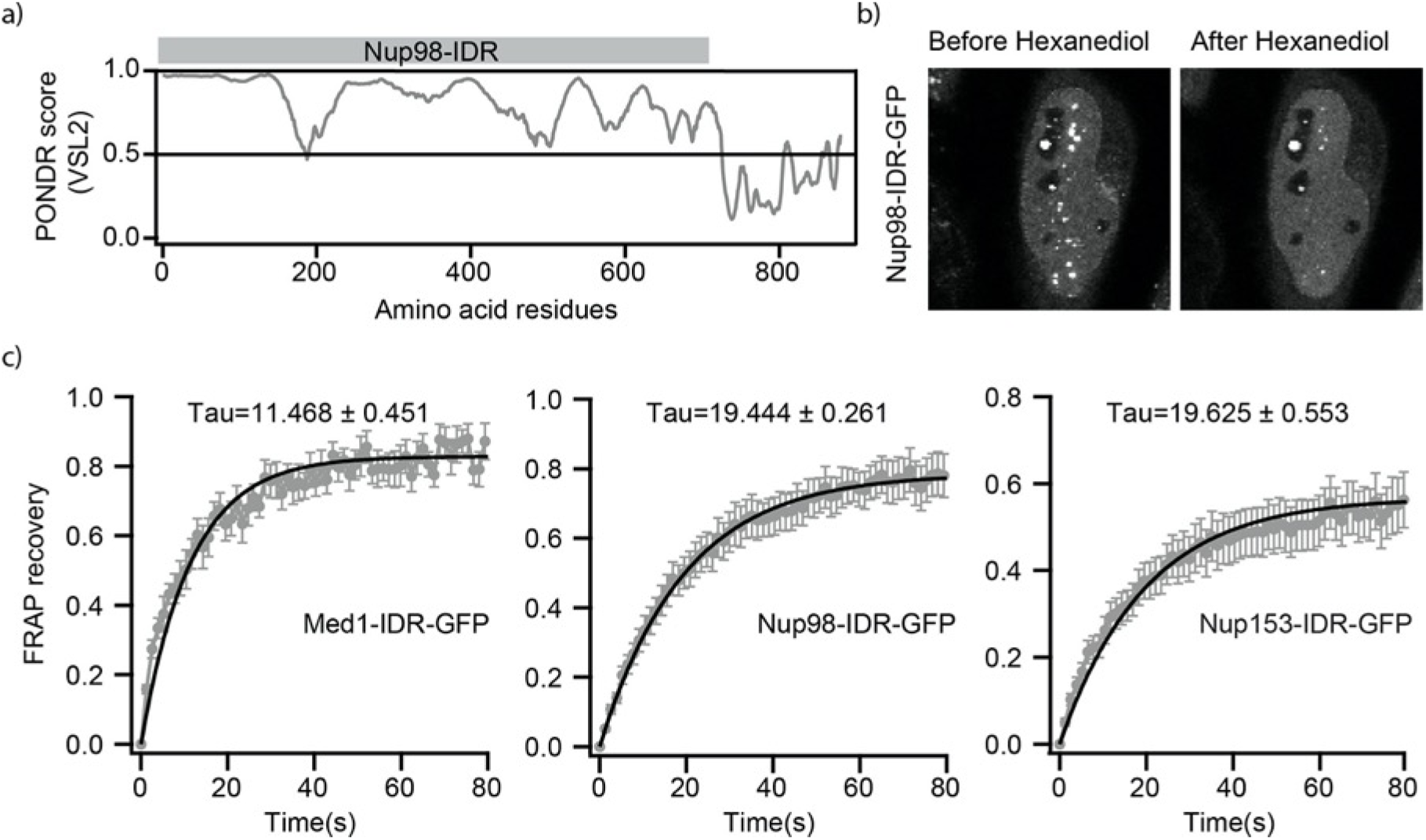
a) PONDR prediction for intrinsically disordered regions (IDR) in the structure of Nup98. A score over 0.5 signifies a lack of structure. b) Confocal images of HeLa cells transfected with GFP fused with Nup98-IDR before and after 30 s treatment with 1.5% hexanediol (HD). c) FRAP recovery plots for 1.5 µm^2^ bleach spot in HeLa nuclei transfected with GFP fused to Med1-IDR, Nup153-IDR, or Nup98-IDR. The solid black line represents the curve fit, and tau is the diffusion time calculated from the curve fits.

**Supplementary Figure 7.**
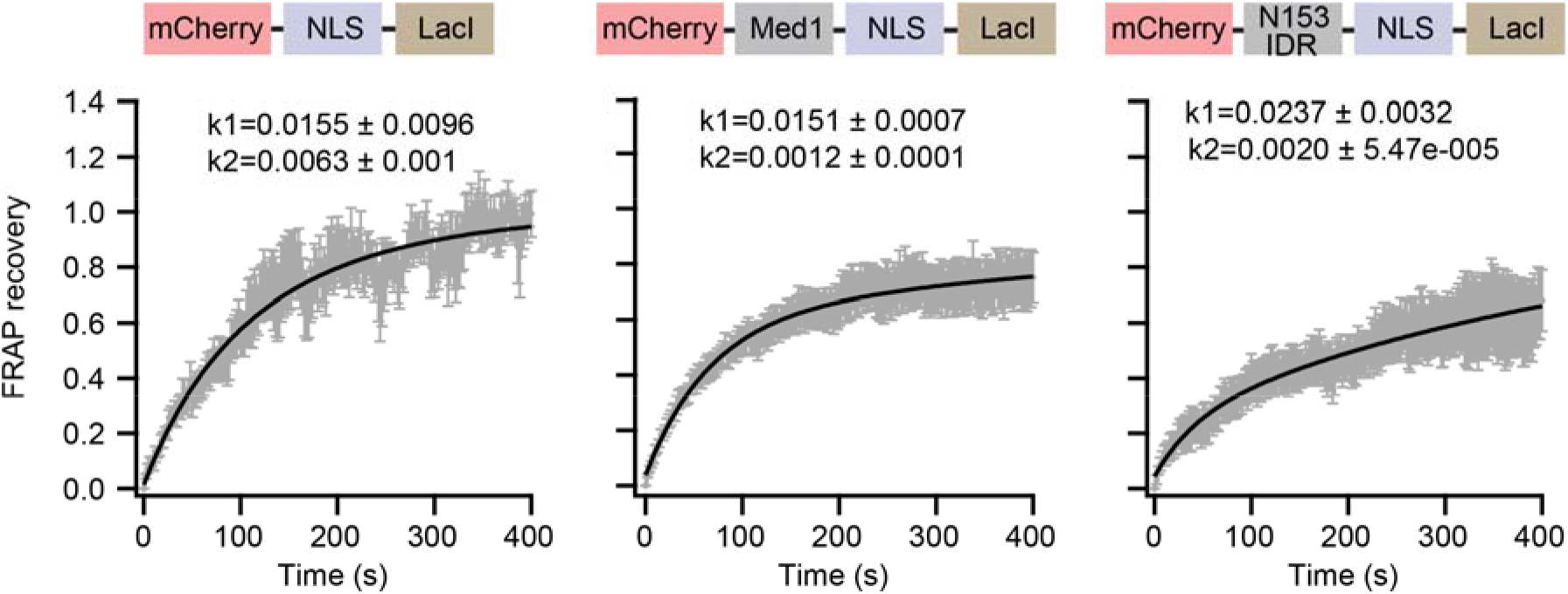
FRAP recovery plots of fluorescent spots for mCherry-NLS-LacI, mCherry-Med1(IDR)-NLS-LacI, and mCherry-Nup153(IDR)-NLS-LacI. The data points are gray, and the solid black line represents the fit. k1 and k2 are dissociationconstants.

**Table S1:** gene annotations around NPC-Dam peaks.

File: TableS1_anno_GSE176106_IMR_DamN44Ibeta_peaks.txt

**Table S2:** coordinates of NPC- and non-NPC SEs and transcription level of their associated genes.

File: TableS2_IMR90.SE.NPCboundornot.FPKMadd.csv

**Table S3:** coordinates of hub and non-hub NPC-SEs.

File: TableS3_enrichment_of_dam_in_hrcySE_hubnonhub.bed

## Notes

http://www.ncbi.nlm.nih.gov/geo/query/acc.cgi?acc=GSE176106

### Competing Interest Statement

The authors have declared no competing interest.

### Summary of Updates

Paper was updated to include additional data and new analysis. New authors contributing to these efforts were added to the paper.

## REFERENCES

1. Cook, P. R. & Marenduzzo, D. Transcription-driven genome organization: a model for chromosome structure and the regulation of gene expression tested through simulations. Nucleic Acids Res 46, gky763 (2018).

2. Lochs, S. J. A., Kefalopoulou, S. & Kind, J. Lamina Associated Domains and Gene Regulation in Development and Cancer. Cells 8, 271 (2019).

3. Ibarra, A. & Hetzer, M. W. Nuclear pore proteins and the control of genome functions. Gene Dev 29, 337–349 (2015).

4. Raices, M. & D’Angelo, M. A. Nuclear Pore Complexes in Genome Organization, Function and Maintenance. 159–182 (2018) doi:10.1007/978-3-319-71614-5_7.

5. D’Angelo, M. A. Nuclear pore complexes as hubs for gene regulation. Nucleus 9, 00–00 (2018).

6. Gatticchi, L., Heras, J. I. de las, Roberti, R. & Schirmer, E. C. Optimization of DamID for use in primary cultures of mouse hepatocytes. Methods 157, 88–99 (2019).

7. Aughey, G. N. & Southall, T. D. Dam it’s good! DamID profiling of protein-DNA interactions. Wiley Interdiscip Rev Dev Biology 5, 25–37 (2016).

8. Aughey, G. N., Cheetham, S. W. & Southall, T. D. DamID as a versatile tool for understanding gene regulation. Development 146, dev173666 (2019).

9. Steensel, B. van & Henikoff, S. Identification of in vivo DNA targets of chromatin proteins using tethered Dam methyltransferase. Nat Biotechnol 18, 424–428 (2000).

10. Southall, T. D. et al. Cell-Type-Specific Profiling of Gene Expression and Chromatin Binding without Cell Isolation: Assaying RNA Pol II Occupancy in Neural Stem Cells. Dev Cell 26, 101–112 (2013).

11. Ibarra, A., Benner, C., Tyagi, S., Cool, J. & Hetzer, M. W. Nucleoporin-mediated regulation of cell identity genes. Gene Dev 30, 2253–2258 (2016).

12. Franks, T. M. et al. Evolution of a transcriptional regulator from a transmembrane nucleoporin. Gene Dev 30, 1155–1171 (2016).

13. Jacinto, F. V., Benner, C. & Hetzer, M. W. The nucleoporin Nup153 regulates embryonic stem cell pluripotency through gene silencing. Gene Dev 29, 1224–38 (2015).

14. Franks, T. M. et al. Nup98 recruits the Wdr82–Set1A/COMPASS complex to promoters to regulate H3K4 trimethylation in hematopoietic progenitor cells. Gene Dev 31, 2222–2234 (2017).

15. Capelson, M. et al. Chromatin-bound nuclear pore components regulate gene expression in higher eukaryotes. Cell 140, 372–83 (2010).

16. Pott, S. & Lieb, J. D. What are super-enhancers? Nat Genet 47, 8–12 (2015).

17. Scholz, B. A. et al. WNT signaling and AHCTF1 promote oncogenic MYC expression through super-enhancer-mediated gene gating. Nat Genet 51, 1723–1731 (2019).

18. Liu, X. et al. In Situ Capture of Chromatin Interactions by Biotinylated dCas9. Cell 170, 1028–1043.e19 (2017).

19. Ryu, J., Kim, H., Yang, D., Lee, A. J. & Jung, I. A new class of constitutively active super-enhancers is associated with fast recovery of 3D chromatin loops. Bmc Bioinformatics 20, 127 (2019).

20. Huang, J. et al. Dissecting super-enhancer hierarchy based on chromatin interactions. Nat Commun 9, 943 (2018).

21. Sabari, B. R. et al. Coactivator condensation at super-enhancers links phase separation and gene control. Science 361, eaar3958 (2018).

22. Mierzejewska, K. et al. Structural basis of the methylation specificity of R.DpnI. Nucleic Acids Res 42, 8745–8754 (2014).

23. Kind, J. et al. Single-Cell Dynamics of Genome-Nuclear Lamina Interactions. Cell 153, 178–192 (2013).

24. Lowe, A. R. et al. Importin-β modulates the permeability of the nuclear pore complex in a Ran-dependent manner. Elife 4, e04052 (2015).

25. Kutay, U., Izaurralde, E., Bischoff, F. R., Mattaj, I. W. & Görlich, D. Dominant-negative mutants of importin-β block multiple pathways of import and export through the nuclear pore complex. Embo J 16, 1153–1163 (1997).

26. Paulsen, J. et al. Chrom3D: three-dimensional genome modeling from Hi-C and nuclear lamin-genome contacts. Genome Biol 18, 21 (2017).

27. Ernst, J. & Kellis, M. Chromatin-state discovery and genome annotation with ChromHMM. Nat Protoc 12, 2478–2492 (2017).

28. Whyte, W. A. et al. Master Transcription Factors and Mediator Establish Super-Enhancers at Key Cell Identity Genes. Cell 153, 307–319 (2013).

29. Xue, B., Dunbrack, R. L., Williams, R. W., Dunker, A. K. & Uversky, V. N. PONDR-FIT: A meta-predictor of intrinsically disordered amino acids. Biochimica Et Biophysica Acta Bba – Proteins Proteom 1804, 996–1010 (2010).

30. Frey, S., Richter, R. P. & Görlich, D. FG-Rich Repeats of Nuclear Pore Proteins Form a Three-Dimensional Meshwork with Hydrogel-Like Properties. Science 314, 815–817 (2006).

31. Labokha, A. A. et al. Systematic analysis of barrier-forming FG hydrogels from Xenopus nuclear pore complexes. Embo J 32, 204–18 (2012).

32. Nott, T. J. et al. Phase Transition of a Disordered Nuage Protein Generates Environmentally Responsive Membraneless Organelles. Mol Cell 57, 936–947 (2015).

33. Chong, S. et al. Imaging dynamic and selective low-complexity domain interactions that control gene transcription, Science 361, eaar2555 (2018).

34. Hung, V. et al. Spatially resolved proteomic mapping in living cells with the engineered peroxidase APEX2. Nat Protoc 11, 456–475 (2016).

35. Ori, A. et al. Cell type-specific nuclear pores: a case in point for context-dependent stoichiometry of molecular machines. Mol Syst Biol 9, 648 (2013).

36. Raices, M. & D’Angelo, M. A. Nuclear pore complex composition: a new regulator of tissue-specific and developmental functions. Nat Rev Mol Cell Bio 13, 687–699 (2012).

37. Gozalo, A. et al. Core Components of the Nuclear Pore Bind Distinct States of Chromatin and Contribute to Polycomb Repression. Mol Cell (2019) doi:10.1016/j.molcel.2019.10.017.

38. Pascual-Garcia, P. et al. Metazoan Nuclear Pores Provide a Scaffold for Poised Genes and Mediate Induced Enhancer-Promoter Contacts. Mol Cell 66, 63–76.e6 (2017).

39. Pascual-Garcia, P. & Capelson, M. Nuclear pores in genome architecture and enhancer function. Curr Opin Cell Biol 58, 126–133 (2019).

40. Kadota, S. et al. Nucleoporin 153 links nuclear pore complex to chromatin architecture by mediating CTCF and cohesin binding. Nat Commun 11, 2606 (2020).

41. Liang, Y., Franks, T. M., Marchetto, M. C., Gage, F. H. & Hetzer, M. W. Dynamic association of NUP98 with the human genome. Plos Genet 9, e1003308 (2013).

42. Light, W. H. et al. A conserved role for human Nup98 in altering chromatin structure and promoting epigenetic transcriptional memory. Plos Biol 11, e1001524 (2013).

43. Iglesias, N. et al. Native Chromatin Proteomics Reveals a Role for Specific Nucleoporins in Heterochromatin Organization and Maintenance. Mol Cell (2019) doi:10.1016/j.molcel.2019.10.018.

44. Labade, A. S., Salvi, A., Karmodiya, K. & Sengupta, K. Nup93 and CTCF co-modulate spatiotemporal dynamics and function of the HOXA gene cluster during differentiation. Biorxiv 6462 24 (2019) doi:10.1101/646224.

45. Sachani, S. S. et al. Nucleoporin 107, 62 and 153 mediate Kcnq1ot1 imprinted domain regulation in extraembryonic endoderm stem cells. Nat Commun 9, 2795 (2018).

46. Blobel, G. Gene gating: a hypothesis. Proc National Acad Sci 82, 8527–8529 (1985).

47. Rajanala, K. et al. Phosphorylation of nucleoporin Tpr governs its differential localization and is required for its mitotic function. J Cell Sci 127, 3505–3520 (2014).

48. McCloskey, A., Ibarra, A. & Hetzer, M. W. Tpr regulates the total number of nuclear pore complexes per cell nucleus. Gene Dev 32, 1321–1331 (2018).

49. Paci, G., Zheng, T., Caria, J., Zilman, A. & Lemke, E. A. Molecular determinants of large cargo transport into the nucleus. Elife 9, e55963 (2020).

50. Milles, S. & Lemke, E. A. Single Molecule Study of the Intrinsically Disordered FG-Repeat Nucleoporin 153. Biophys J 101, 1710–1719 (2011).

51. Marshall, O. J., Southall, T. D., Cheetham, S. W. & Brand, A. H. Cell-type-specific profiling of protein–DNA interactions without cell isolation using targeted DamID with next-generation sequencing. Nat Protoc 11, 1586–1598 (2016).

52. Chen, C. et al. SEA version 3.0: a comprehensive extension and update of the Super-Enhancer archive. Nucleic Acids Res 48, D198–D203 (2019).

